# Aβ receptors specifically recognize molecular features displayed by fibril ends and neurotoxic oligomers

**DOI:** 10.1101/822361

**Authors:** Ladan Amin, David A. Harris

## Abstract

Oligomeric forms of amyloid-β (Aβ) peptide are known to be the primary neurotoxic species in Alzheimer’s disease (AD), but how they interact with neurons to produce their deleterious effects is unclear. Over ten different cell-surface receptors for Aβ have been described, but their molecular interactions with Aβ assemblies and their relative contributions to mediating AD pathology have remained uncertain. In the present work, we have used super-resolution microscopy to directly visualize Aβ-receptor interactions at the nanometer scale. We report that one documented Aβ receptor, the cellular prion protein, PrP^C^, specifically inhibits the polymerization Aβ fibrils via a unique mechanism in which it binds specifically to the rapidly growing end of each fibril, thereby blocking polarized elongation at that end. PrP^C^ binds neurotoxic oligomers and protofibrils in a similar fashion, suggesting that it may recognize a common, end-specific, structural motif on all of these assemblies. Finally, two other candidate Aβ receptors, FcγRIIb and LilrB2, affect Aβ fibril growth in a manner similar to PrP^C^. Taken together, our results suggest that neurotoxic signaling by several different receptors may be activated by common molecular interactions with both fibrillar and oligomeric Aβ ligands. Targeting such interactions with small molecules represents an attractive therapeutic strategy for treatment of AD.

## INTRODUCTION

Alzheimer’s disease (AD) is a progressive neurodegenerative disease that is characterized by accumulation within the brain of extracellular plaques composed of the amyloid-β (Aβ) peptide, and intracellular neurofibrillary tangles containing abnormally phosphorylated forms of the tau protein^1,2^. These two kinds of pathological deposit lead ultimately to synaptic loss and dysfunction, and to degeneration and loss of neurons. Aβ peptides of 40-42 amino acids in length are derived from the amyloid precursor protein (APP) via sequential cleavage by the enzymes β- and γ- secretase^2,3^. In AD brain, Aβ is either over-produced and/or degraded inefficiently, resulting in the formation of several types of Aβ aggregate^4^. There is strong evidence that small Aβ oligomers (Aβo), rather than monomers or fibrils, represent the key neurotoxic species in AD^5–7^. It is presumed that the disease process starts by the binding of Aβo to receptor proteins or lipids on the surface of neurons. However, the molecular identity of the relevant binding sites, and the signal transduction pathways they trigger leading to synaptotoxicity, are uncertain. Identification of Aβ receptors and elucidation of their mechanism of interaction with Aβo have important therapeutic implications, since these receptors represent potential pharmacological targets for treatment of AD.

A number of cell-surface proteins have been reported to act as Aβ receptors^8,9^. Among them, the cellular prion protein (PrP^C^)^10^, Fcγ receptor IIb (FcγRIIb)^11^, leukocyte immunoglobulin-like receptor; subfamily B2 (LilrB2)^12^, ephrin type-B receptor 2 (EphB2)^13^, and Nogo receptor family (Ngr1-3)^14^, have attracted particular attention because of their high affinity for Aβo, and their ability to transduce a neurotoxic signal. However, there has been considerable controversy about the relative importance of each of these receptors in mediating Aβ neurotoxicity, with discrepant results emanating from many of the published studies^15–18^. At this point, it seems reasonable to assume that there are multiple receptors capable of binding Aβ assemblies in a physiological context and mediating their synaptotoxic actions. Although interaction of Aβ with each of the receptors has previously been shown using *in vitro* or cellular binding assays, the molecular details of the binding reaction remain unclear.

Recently, PrP^C^ was identified as a high-affinity receptor for Aβo^10^ an observation subsequently confirmed by other groups^19–23^. Physical binding of Aβo and PrP^C^ has been demonstrated using both cellular and biochemical methods^10,19–23^. Binding of Aβo to PrP^C^ has also been reported to initiate synaptotoxic signaling, causing suppression of long-term potentiation (LTP) and retraction of dendritic spines^10,24,25^. It has been proposed that this signaling pathway requires interactions between PrP^C^ and mGluR5, resulting in activation of intracellular fyn kinase, with subsequent phosphorylation and redistribution of NMDA receptors^25–28^. In *vivo*, genetic deletion of PrP^C^ rescues behavioral deficits as well as early mortality in certain AD transgenic models^29^. Compounds that block Aβ binding to PrP^C^, or that inhibit activation of downstream signaling mechanisms, have been shown to ameliorate pathology in AD transgenic models^30,31^, and some of these are being tested in human patients^32^.

Amyloid fibril formation from soluble Aβ monomers is a well-characterized process that involves distinct, kinetically defined steps of primary nucleation, secondary nucleation and elongation^33–35^. Several classes of molecules, including chaperones, antibodies, and small molecules, have been reported to influence specific steps of the polymerization process^34,36–39^. Our laboratory has recently investigated the effect of PrP^C^ on Aβ polymerization^22^. That study, which relied upon biochemical assays and mathematical modeling, demonstrated that PrP^C^ specifically inhibits the elongation step of Aβ polymerization, most likely by binding to the ends of growing fibrils. However, we did not directly prove this mechanism by measurement of fibril elongation rates, or by localization PrP on individual fibrils.

In this study, we have employed super-resolution microscopy to directly visualize, at a nanoscale level, the dynamics of Aβ assembly, and the interactions of PrP^C^, as well as two other cell surface receptors, with Aβ fibrils and neurotoxic oligomers. Analyzing several Aβ-receptor systems in parallel enabled us to reveal common molecular mechanisms by which these receptors interact with pathologically relevant Aβ aggregates to transduce neurotoxic signals. Altogether, our data provide new insights into the molecular origins of AD, and they lay the groundwork for development of therapeutic approaches to block receptor-mediated Aβ neurotoxicity.

## RESULTS

In this study, we have used direct stochastic optical reconstruction microscopy (dSTORM)^40^ and structured illumination microscopy (SIM)^41^ to visualize directly the effect of PrP and two other putative Aβ receptors on the process of Aβ polymerization with a resolution of 20 nm (dSTORM) or 100 nm (SIM) (Supplementary Fig. S1). At these resolutions, we were able to visualize Aβ oligomers, protofibrils and fibrils, and determine the localization of PrP on these structures. We polymerized synthetic Aβ1-42 labeled with either Cy3 or Cy5 under carefully defined conditions, which have been shown to result in reproducible kinetic curves as monitored by thioflavin T (ThT) fluorescence^33,34,42^. For co-localization experiments, recombinant PrP was fluorescently labeled by substituting a cysteine residue at position 34, and reacting it with a maleimide derivative of either Alexa Fluor 555 (AF555) or Alexa Fluor 488 (AF488). We first monitored the kinetics of fibril formation by ThT fluorescence to confirm that incorporation of the fluorescent labels into either Aβ or PrP did not affect the Aβ polymerization process or the ability of PrP to inhibit polymerization (Supplementary Fig. S2). As we reported previously^22^, PrP in sub-stoichiometric amounts dramatically inhibited Aβ polymerization.

### PrP promotes formation of shorter, more numerous fibrils

Based on its effect on the kinetics of Aβ polymerization, we previously concluded that PrP specifically inhibited the elongation step of polymerization^22^. In this case, it would be predicted that Aβ fibrils formed in the presence of PrP would be shorter than those formed in the absence of PrP. Moreover, the total number of fibrils would be increased in the presence of PrP, since the flux of monomers would be shifted from elongation toward nucleation events that generate additional fibrils^37^. To test these predictions, Aβ-Cy5 monomers at a concentration of 20 μM were polymerized for 24 h in the presence of different concentrations of PrP. The fibrils that formed were then imaged by SIM.

We found that PrP over a concentration range of 0.1-1 μM caused a dose-dependent reduction in the length of Aβ fibrils, and significantly increased the number of fibrils (Fig. 1A-D). To quantify these effects, we measured the length and number of all the fibrils resolved in each SIM image. We found that, when the PrP concentration was increased from 0 to 1 μM, the mean value of fibril length decreased progressively from 0.90±0.02 μm to 0.29±0.01 μm (Fig. 1E), while the number of fibrils increased from 0.13±0.005/μm^2^ to 0.66±0.02/μm^2^ (Fig. 1G). Although these preparations were heterogeneous, the size distribution of fibrils formed in presence of PrP was significantly different from control samples, with a preponderance of smaller species in the former samples (Fig. 1F, compare red curve with grey curves). In the absence of PrP, some fibrils reached lengths of up to 6 μm, and approximately 10% of the fibrils were >2 μm (Fig. 1F, inset). In contrast, even at lowest concentration of PrP (0.1 μM), no fibrils were longer than 3 μm, and only 2.7% were >2 μm. Increasing the concentration of PrP shifted the population of Aβ fibrils to smaller sizes.

**Fig. 1.**
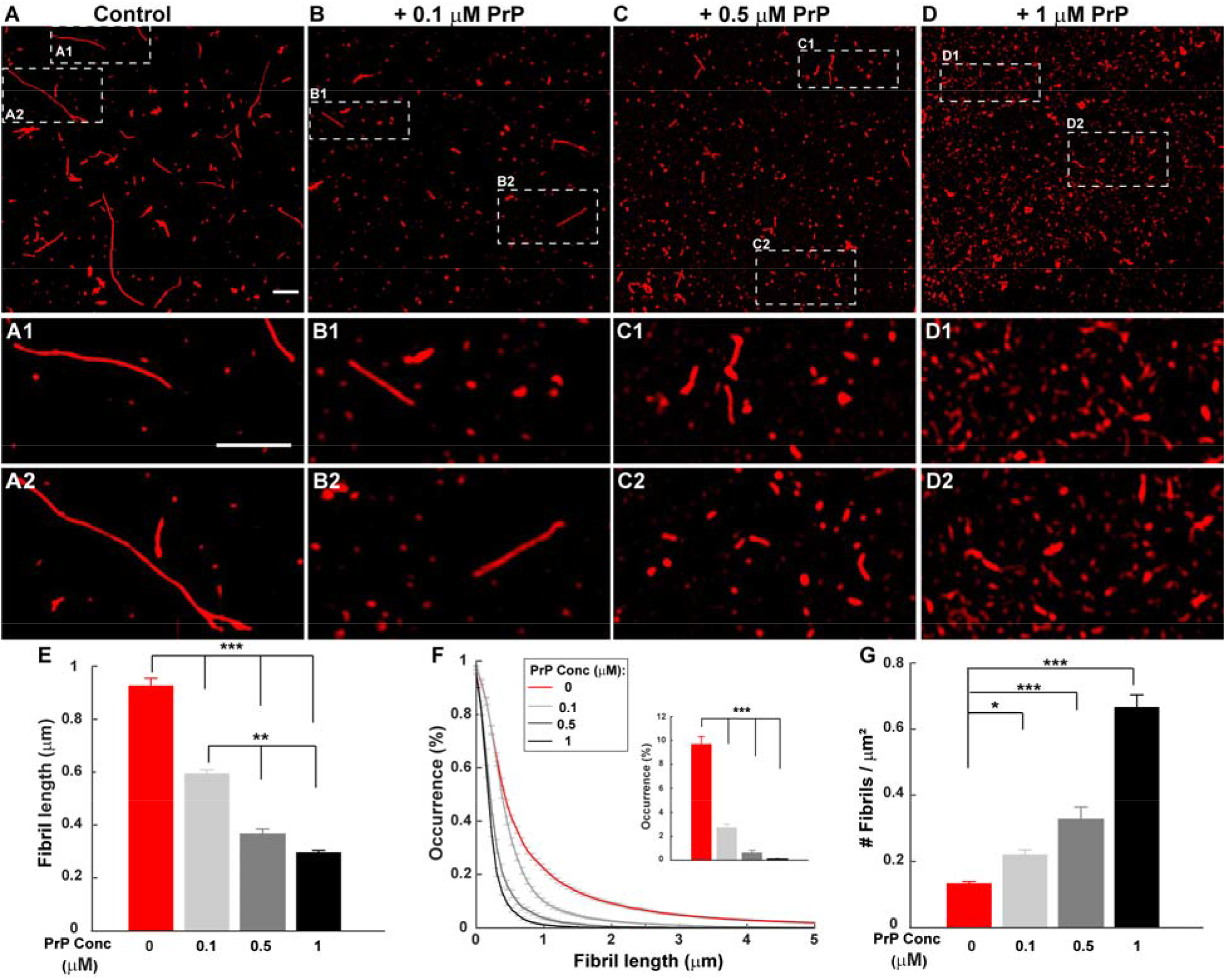
PrP promotes formation of shorter, more numerous Aβ fibrils. Aβ-Cy5 monomer (20 μM) was polymerized for 24 h in the presence of 0 μM **(A)**, 0.1 μM **(B)**, 0.5 μM **(C)**, or 1 μM **(D)** PrP-AF555. Fibrils were then imaged by SIM. Panels A1,2-D1,2 show boxed areas in A-D, respectively, at higher magnification. Scale bars are 1 μm. **(E)** Bars show mean fibril length at each PrP concentration. **(F)** Cumulative distributions of fibril length at each PrP concentration. Inset indicates the number of fibrils larger than 2 μm. **(G)** Bars indicate the number of detectable Aβ-Cy5 fibrils/μm^2^ at each PrP concentration. Data represent mean ± S.E. * P<0.05, ** P<0.01 and *** P < 0.001 (Student’s t-test).

Taken together, these data suggest that PrP affects Aβ polymerization in a dose-dependent manner, resulting in the formation of shorter but more numerous fibrils at 24 h.

### PrP slows the growth of Aβ fibrils

To characterize directly the effect of PrP on the dynamics of the elongation process, we measured the effect of PrP on fibril lengths at different points of the Aβ polymerization reaction. Each assay began with 20 μM Aβ in monomeric form. In the absence of PrP, Aβ monomers rapidly polymerized, reaching a maximum size distribution by 24 h, after which mean fibril size remained constant for up to seven days (Fig. 2A; quantitation in Fig. 2C-D). In contrast, the polymerization rate was slowed significantly when 0.5 μM PrP-AF555 was added to the reaction at the starting point. Under these conditions, the fibrils continued to grow slowly over seven days (Fig. 2B; quantitation in Fig. 2C-D). At each of the time points analyzed, the mean length of fibrils formed in presence of PrP was significantly less than in control conditions. In addition, the number of fibrils was higher in the presence of PrP at each time point, from 4 hrs to 7 days, after the initiation of polymerization (Fig. 2E). Taken together, these results indicate that PrP significantly slows the process of fibril elongation, and increases the number of fibrils formed.

**Fig. 2.**
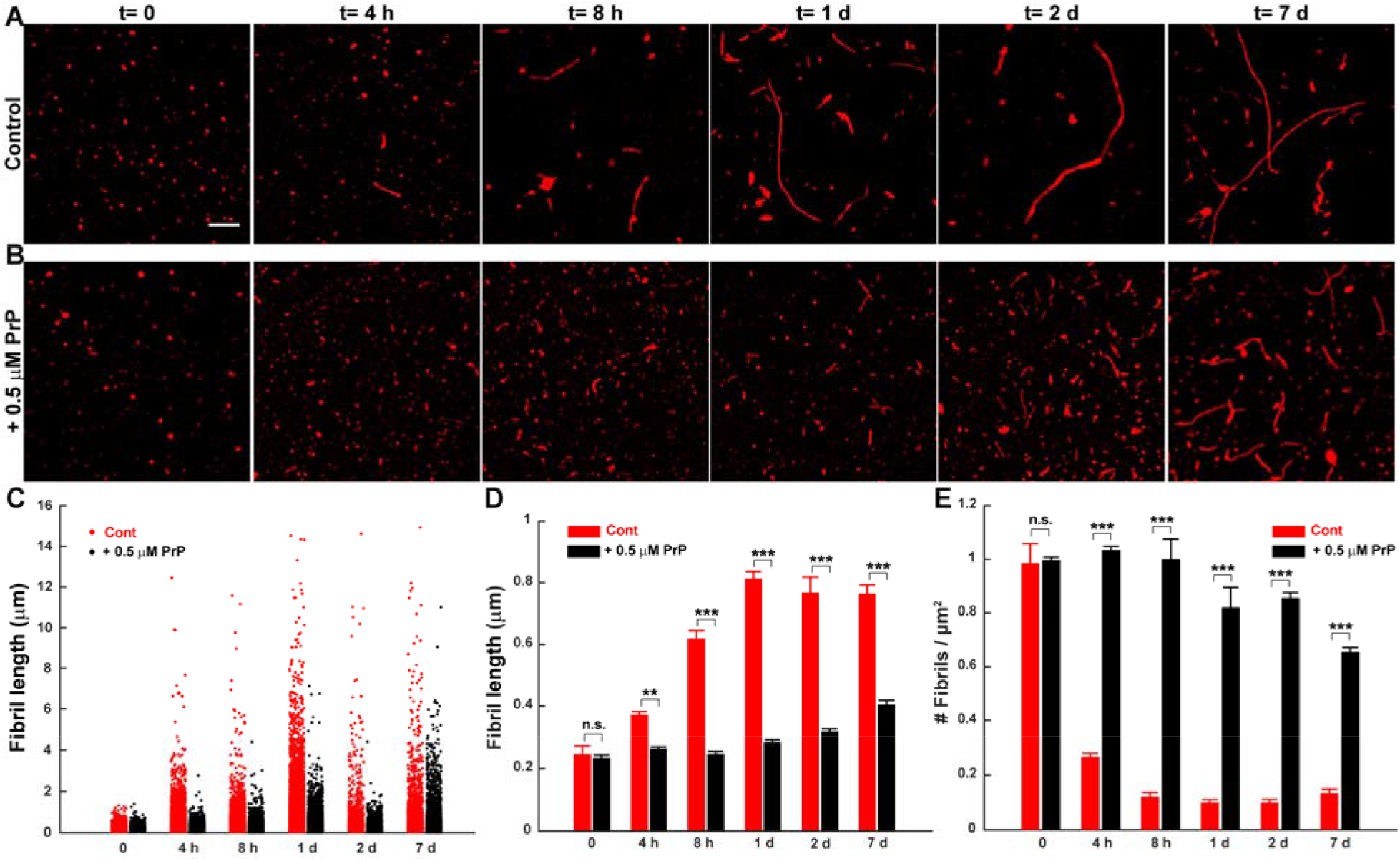
PrP slows the growth of Aβ fibrils. Aβ-Cy5 monomer (20 μM) was polymerized for the indicated times in the presence of 0 μM **(A)** or 0.5 μM **(B)** PrP-AF555. Fibrils were then imaged by SIM. Scale bar in panel A (t=0) is 1 μm. **(C)** Distributions of fibril lengths at each time point in the presence of 0 μm (red dots) or 0.5 μM PrP-AF555 (black dots). Each dot represents an individual fibril. **(D, E)** Bars indicate the mean length of fibrils **(D)** and the number of detectable Aβ-Cy5 fibrils/μm^2^ **(E)** at each time point for 0 and 0.5 μM PrP-AF555. Data represent mean ± S.E. * P<0.05, ** P<0.01 and *** P < 0.001 (Student’s t-test).

### Aβ polymerization is strongly polarized, and PrP selectively blocks elongation at the more rapidly growing end

It has been shown that elongation of Aβ fibrils is strongly polarized, with the two ends of the fibril growing at different rates (fast and slow)^43^. By adapting several published seeding procedures^44–46^, we were able to determine how PrP affected the elongation rate at each end of the fibril. Sheared, preformed fibrils (referred to as seeds) labeled with Cy5 were allowed to grow by incubation in a solution of 10 μM monomeric Aβ labeled with Cy3 (Fig. 3A). After different lengths of time, ranging from 2 hrs to 7 days, the fibrils were imaged by SIM to visualize growth of the seeds at their ends. The lengths of the green extensions at the two ends of each red seed were measured over time to monitor the progress of elongation.

**Fig. 3.**
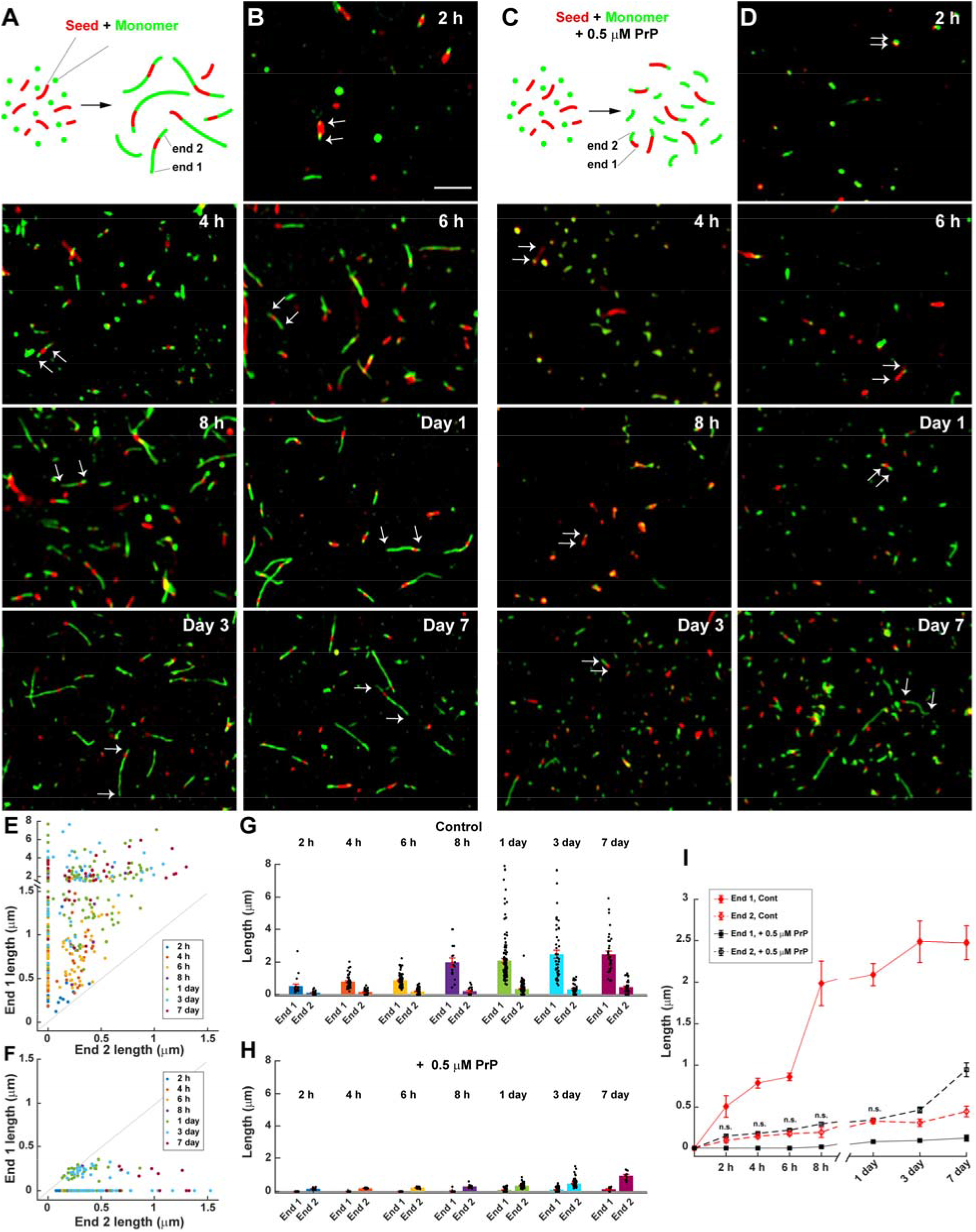
Aβ polymerization is strongly polarized, and PrP selectively blocks elongation at the fast growing end. **(A)** Schematic representation of a seeding assay, in which fresh monomers labeled with Cy3 (green) were added to sheared, preformed fibrils (seeds) labeled with Cy5 (red). The two ends of each seed elongate at different rates, resulting in long and short green extensions, designated End 1 and End 2, respectively. **(B)** Two-color SIM images acquired at the indicated times after addition of Aβ-Cy3 monomers (10 μM) to Aβ-Cy5 seeds (10 μM monomer equivalent). Arrows in each panel indicate the elongation of the seed at the two ends. Scale bar in panel A (2 hr) is 2 μm. **(C)** Schematic representation of the seeding assay in the presence of 0.5 μM PrP, indicating that seeds elongate only at End 2. **(D)** Two-color SIM images acquired as in Panel B, but in the presence of 0.5 μM PrP. Arrows in each panel show that seeds elongate at only one end. **(E, F)** Scatterplots showing the lengths of End 1 and End 2 over time for each detected seed in the absence of PrP (E) and in the presence of 0.5 μM PrP (F). The total number of seeds measured was 390 and 215 for 0 μM and 0.5 μM PrP, respectively. **(G, H)** Bars indicate the mean lengths of End 1 and End 2 over time in the absence of PrP (G) and in the presence of 0.5 μM PrP (H). Data represent mean ± S.E. **(I)** Change in the mean lengths of End 1 and End 2 over time, with and without PrP. These are the same data as in panels F and G, but plotted to allow easier comparison of the different conditions. n.s., no statistically significant difference between End 2 length with and without PrP at the indicated time points.

Using this assay, we confirmed a previous report^43^ that fibril growth under control conditions occurs asymmetrically at the two ends (Fig. 3A and B). At each time point, one end of every fibril had grown more than the other end, as revealed by a scatterplot of the lengths of the green extensions at the two ends of 390 seeds (Fig. 3E). There was no statistical correlation between the lengths of the extensions at the two ends of each fibril, indicating that the two ends behaved independently. We noted that some seeds grew only at one end during the course of the experiment, a phenomenon that was observed previously, and was attributed to longer paused periods in fibril growth at the slow end^43^. We have designated the longer (faster-growing) extension as End 1, and the shorter (slower-growing) extension as End 2. At early time points (2 h), the mean lengths of the fast- and slow-growing ends were 0.50±0.13 and 0.11±0.02 μm, respectively (Fig. 3G and I). By 24 hrs, the mean lengths of the fast and slow-growing ends had reached 2.09 ± 0.13 μm and 0.32 ± 0.03 μm, respectively, after which they changed relatively little for up to seven days (Fig. 3G and I).

When 0.5 μM PrP was added to the reaction together with fresh monomers, the growth characteristics were markedly different (Fig. 3C and D). Green extensions were seen at only one end of most fibrils (Fig. 3F), and these extensions grew slowly, with mean lengths of 0.14±0.017 at 2 h, 0.34 ± 0.017 at 1 day, and 0.94 ± 0.08 at 7 days (Fig. 3H and I). The elongation values at all but the seven-day time point were statistically indistinguishable from those measured at the slow-growing end in the absence of PrP (Fig. 3I, compare black and red dashed lines). Taken together, these data suggest that PrP completely blocks elongation at the fast-growing end of the fibril, with growth of the fibril restricted to elongation at the slow-growing end.

### PrP binds exclusively to the fast-growing end of Aβ fibrils

A likely mechanism by which PrP blocks fibril elongation is by binding selectively to the fast-growing end of the fibril, preventing further monomer addition at that end. To directly localize PrP on individual fibrils, we polymerized Aβ-Cy5 monomers in the presence of different concentrations of PrP-AF555 for 24 h, and then visualized Aβ aggregates by super-resolution microscopy. Strikingly, dual-color dSTORM images showed that PrP-AF555 was selectively associated with only one end of each Aβ fibril (Fig. 4A). As would be expected, the number of aggregates with co-localized PrP increased as the concentration of PrP was raised (Fig. 4B).

**Fig. 4.**
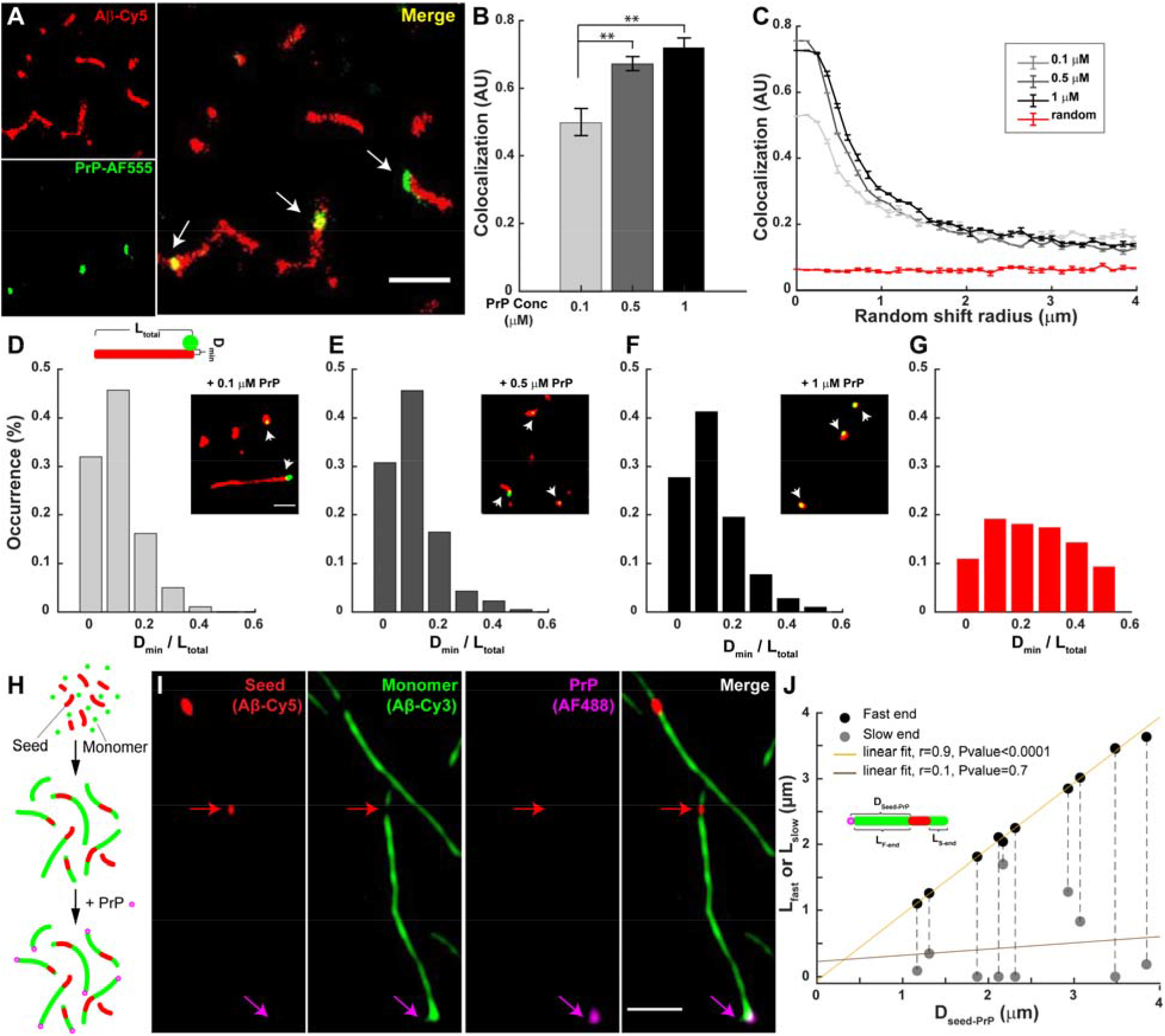
PrP binds exclusively to the fast-growing end of Aβ fibrils. **(A)** dSTORM images of Aβ-Cy5 (20 μM, red) polymerized for 24 h in the presence of 0.5 μM PrP-AF555 (green). Arrows in the merged image indicate the localization of PrP near the ends of Aβ fibrils. Scale bar is 1 μm. **(B)** Bars indicate the colocalization (calculated as described in Methods) between Aβ and PrP-AF555 at three different concentration of PrP. A.U., arbitrary units. **(C)** Colocalization between Aβ and PrP decays after a set of random direction shifts, and approaches the values derived from unrelated PrP and Aβ images from two different experiments (red line). **(D-F)** Aβ-Cy5 (20 μM) was polymerized for 24 h in the presence of either 0.1 μM, 0.5 μM, or 1 μM PrP-AF555, and samples were imaged by dSTORM. The minimum distance (D_min_) between the PrP spot and the closest end of the underlying fibril was measured, and normalized to the total length (L_total_) of the fibril, to give the quantity D_min_/L_total_ (see cartoon in panel D). Histograms show the distribution of D_min_/L_total_ values for 95-227 fibrils with associated PrP spots. Insets show the dSTORM images of Aβ fibrils (red) formed in the present of different concentration of PrP-AF555 (green). Arrowheads show the position of PrP-AF4555 on individual fibrils. Scale bar in panel D inset is 0.5 μm. **(G)** Random distribution of D_min_/L_total_ values derived from unrelated PrP and Aβ images from two different experiments. **(H)** Schematic representation of three-color imaging assay. Preformed seeds composed of Aβ-Cy5 (red) were first incubated for 24 hrs with monomeric Aβ-Cy3 (green), to allow extension of the seeds at both ends. PrP-AF488 (magenta) was then added for 30 min, and the fibrils were imaged by three-color SIM. **(I)** Individual fibrils imaged for Cy5 (Aβ seed), Cy3 (Aβ monomer), AF488 (PrP), and a merge of the three colors. The magenta arrow indicates PrP bound to the fast-growing end of a single fibril, represented by the long green extension from a red seed. The red arrow indicates the position of the seed. Scale bar in the merged panel is 1 μm. **(J)** Distance between the seed and the PrP spot (D_seed-PrP_) plotted against the lengths of the fast-and slow-growing ends (L_fast_ and L_slow_, respectively) for 10 separate fibrils (see cartoon for definitions of these measurements). The pairs of black and grey dots connected by a dotted line correspond to the two ends of each fibril. D_seed-PrP_ is the same as L_fast_, reflecting the fact that the PrP is always bound at the end of the long (fast-growing) extension. See Supplementary Fig. S3 for additional images of PrP bound to the fast-growing end of the fibril.

To demonstrate that the observed association of PrP with Aβ aggregates was specific, and not the result of random colocalization, we employed a randomization test. First, the position of PrP in each image was digitally shifted in a random direction along a two-dimensional vector of defined size, and then the colocalization index was recomputed. We performed this analysis for different vector sizes ranging from 100 nm to 4 μm (Fig. 4C). We found that the colocalization index decreased rapidly as the size of the vector was increased, reaching a minimum level, corresponding to random placement. As a control, we measured the colocalization between PrP and Aβ in unrelated images from two different experiments (red curve in Fig. 4C). In this case, very low colocalization was measured, and this value remained constant when the size of the vector increased. Taken together, these results demonstrate that PrP associates non-randomly with individual Aβ aggregates.

We next adopted an unbiased statistical method to quantify the localization of PrP with respect to fibril ends for large populations of individual Aβ fibrils over a range of different PrP concentrations. We measured the minimum distance (D_min_) between each fluorescent PrP spot and the closest end of the associated fibril, and then normalized this distance to the total length (Ltotal) of the fibril to give the quantity D_min_/L_total_ (see cartoon in Fig. 4D). D_min_/L_total_ values could, theoretically, vary between 0 (PrP exactly at the fibril end) and 0.5 (PrP at the mid-point of the fibril). We chose this normalization procedure, since fibril length decreases significantly with increasing amounts of PrP (Figs. 1 and 2). When we plotted the distribution of the D_min_/L_total_ values for a large number of fibrils, we found that, at each of three different PrP concentrations, the majority of D_min_/L_total_ values were less than 0.1, indicating that the PrP spot was localized very close to one end of the underlying fibril (Fig. 4D-F). At 0.1, 0.5, and 1 μM PrP, the proportion of D_min_/L_total_ values <0.1 was 77%, 75%, and 68%, respectively. Thus, even for the very short fibrils present at high PrP concentrations, the PrP spot was localized asymmetrically, closer to one end of the fibril. At all three tested concentrations, the localization of PrP on Aβ fibrils was significantly different from a random distribution, which was determined by measuring D_min_/L_total_ distances in images from unrelated PrP and Aβ experiments (Fig. 4G). In this case, only 30% of the D_min_/L_total_ values were less than 0.1.

We next wished to determine whether the fibril end where PrP bound was the fast- or slow-growing end. To address to this question, we took advantage of our seeding assay, in which we could clearly resolve fast and slow growing ends of individual fibrils (Fig. 4H). In these experiments, fibrils were polymerized by addition of 10 μM Aβ-Cy3 monomer (green) to short, preformed seeds consisting of Aβ-Cy5 (red). After 24 h of incubation at 37°C, PrP-AF488 (magenta) was added to seeded fibrils, and samples were then imaged by three-color SIM. We found that PrP-AF488 was selectively associated with the end of the fibril with the longer green extension, indicating binding to the fast-growing end of the fibril (Fig. 4I and Supplementary Fig. S3). This localization was consistent over a range of different lengths of the extensions from the fast-growing end, and was not correlated with the lengths of the extensions from the slow-growing end (Fig. 4J). These data indicate that PrP inhibits fibril elongation by binding selectively to the fast-growing end of the fibril, thereby blocking growth at that end.

### Localization of PrP on neurotoxic Aβ assemblies

Our results thus far have focused on the localization of PrP on Aβ fibrils. However, small oligomeric assemblies of Aβ, rather than long fibrils, are thought to be the most neurotoxic species, and to be primarily responsible for synaptic loss and neurological dysfunction in AD^5–7^. We therefore sought to define the localization of PrP on two kinds of neurotoxic Aβ assemblies: Aβ-derived diffusible ligands^47^ (ADDLs) and protofibrils^48^. These two kinds of Aβ assemblies, although differing in their method of preparation and structural properties, share the common feature of being highly neurotoxic when tested in cellular, brain slice, and *in vivo* assays. ADDLs represent a heterogeneous collection of oligomeric species, which typically appear in EM images as globular or ellipsoid structures of approximately 20 nm diameter^47^, near the resolution limit of dSTORM. Protofibrils (formed by incubation of ADDLs for up to 1 week) are larger aggregates that have the appearance of short fibrils of 20-200 nm in length^21^. For comparison, we also localized PrP on fully polymerized Aβ fibrils.

In the first set of experiments, ADDLs, Aβ protofibrils and fibrils were prepared using Cy5-labeled peptide, and were then incubated with PrP-AF555. Samples were then imaged with dual-color dSTORM. As expected, PrP and ADDLs co-localized with a high degree of overlap (Fig. 5A). Interestingly, we often observed that PrP localized eccentrically on globular or ellipsoid-shaped ADDL aggregates, being closer to one edge of the aggregate (Fig 5A1). This effect was apparent when we calculated D_min_/L_total_ values for a large number of aggregates; these values were not evenly distributed, with 82% of the aggregates displaying D_min_/L_total_ values <0.1, indicative of an asymmetric localization (Fig. 5B). On protofibrils, which typically displayed identifiable ends, we observed that PrP was bound exclusively to one end (Fig. 5C and C1), similar to its localization on longer Aβ fibrils (Fig. 5E and E1). On protofibrils and fibrils, 56% and 73% of the D_min_/L_total_ values, respectively, were <0.1 (Fig. 5D and F)

**Fig. 5.**
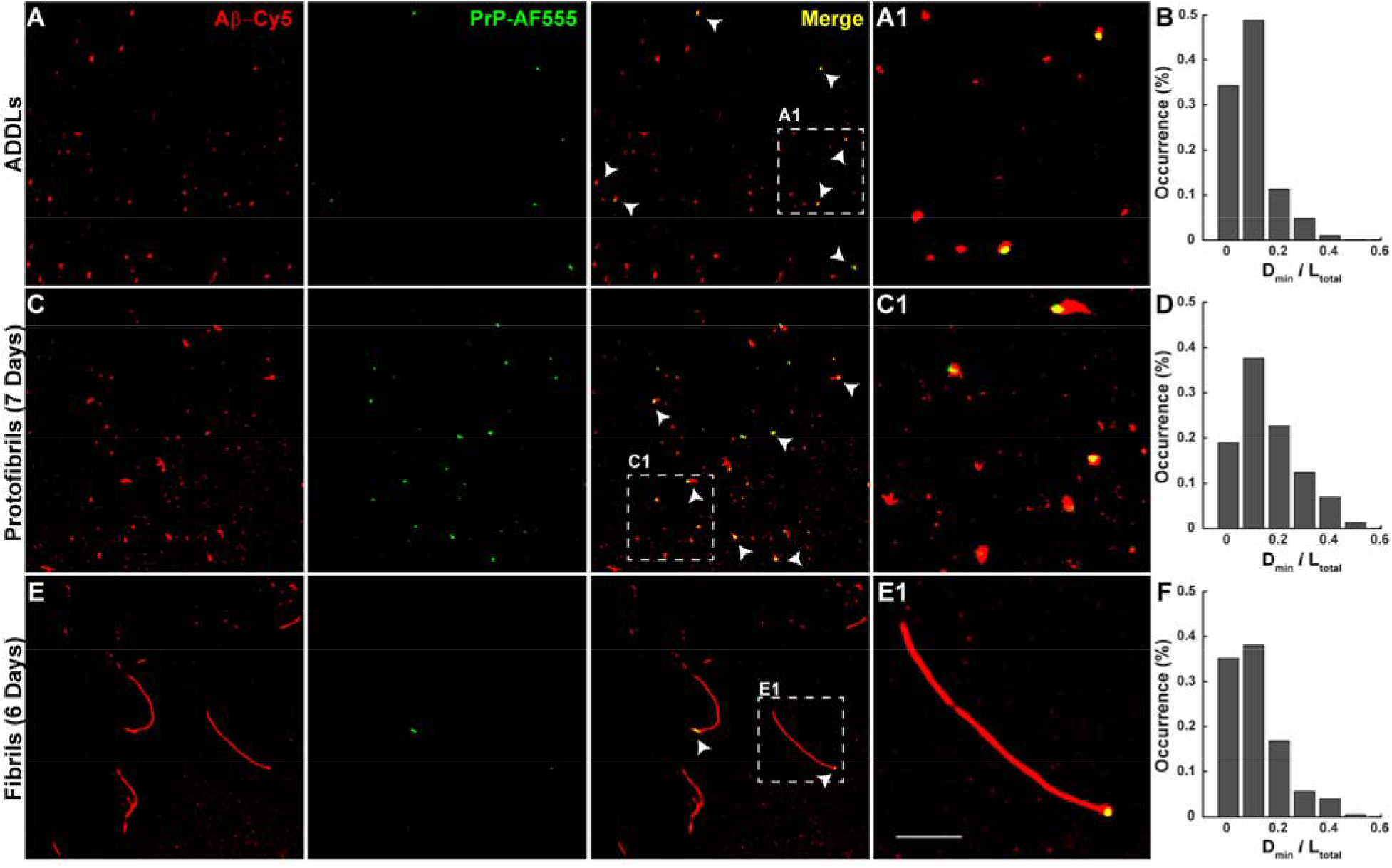
Localization of PrP on neurotoxic Aβ assemblies. Pre-formed ADDLs (**A**), protofibrils (**C**), and fibrils (**E**) (all at 20 μM monomer-equivalent concentration) were incubated with 0.5 μM PrP-AF555 and then imaged by dSTORM. Panels A1, C1, and E1 show boxed areas in A, C and E, respectively, at higher magnification. Scale bar in panel E1 is 1 μm. Histograms (**B, D, F**) show the distribution of D_min_/L_total_ values, calculated as in Fig. 4, for ADDLs, protofibrils, and fibrils, respectively.

In a second set of experiments, we determined the effect of PrP on protofibril formation. PrP was added to ADDL preparations, and samples were then incubated for one week to allow the formation of protofibrils. We found that protofibrils formed in the presence of PrP were significantly shorter than species formed in control samples, and PrP again bound exclusively to one end of each protofibril (Supplementary Fig. S4). Taken together, these data suggest that PrP interacts selectively with oligomer and protofibril ends in a fashion similar to its interaction with the ends of longer Aβ fibrils.

We assessed the neurotoxicity of each of Aβ aggregates used in these experiments by assaying their ability to induce retraction of dendritic spines on cultured hippocampal neurons^49^ (Supplementary Fig. S5). We have shown previously that Aβ oligomer toxicity in this assay is PrP^C^-dependent^50^. We found that toxicity was correlated with the size of aggregate; ADDLs and 3-day protofibrils induced more marked spine retraction than 7-day protofibrils, while fully polymerized fibrils were relatively inert. These data confirm previous evidence that smaller oligomers are more toxic than fibrils^5^, and they demonstrate that the ADDL and protofibril preparations used for our experiments are indeed neurotoxic in a biologically relevant assay.

### Other putative receptors interact with Aβ in a manner similar to PrP^C^

Having established a model for Aβ-PrP^C^ interactions, we asked whether other putative receptor proteins interacted with Aβ aggregates via a similar mechanism. We decided to focus on FcγRIIb and LilrB2, since there is strong evidence that these receptors bind Aβ oligomers, and transduce neurotoxic signals in biological assays^11,12,51^. FcγRIIb, which is expressed in B cells, macrophages, neutrophils, as well as neurons, binds antigen-bound IgG complexes and transduces an inhibitory signal that results in inhibition of the B-cell-mediated immune response^52^. LilrB2 (whose mouse ortholog is called PirB) was originally thought to function exclusively in the immune system, but is now known to be expressed by neurons and to be involved in neurodevelopmental events^53^.

For these experiments, we used the purified, extracellular domains of FcγRIIb and LilrB2, which encompass the known Aβ binding sites^11,12^. First, we analyzed the effect of each of these receptor proteins on Aβ polymerization kinetics using ThT fluorescence (Fig. 6). We found that both FcγRIIb and LilrB2 inhibited Aβ polymerization in a manner very similar to PrP^C^ (Fig. 6A-C), while a control protein, calmodulin, had no effect (Fig. 6D). In sub-stoichiometric amounts, each of these receptor proteins increased the half-time required to reach a plateau value of ThT fluorescence (Fig. 6E). In contrast, the half-time remained constant for even very high concentrations of calmodulin.

**Fig. 6.**
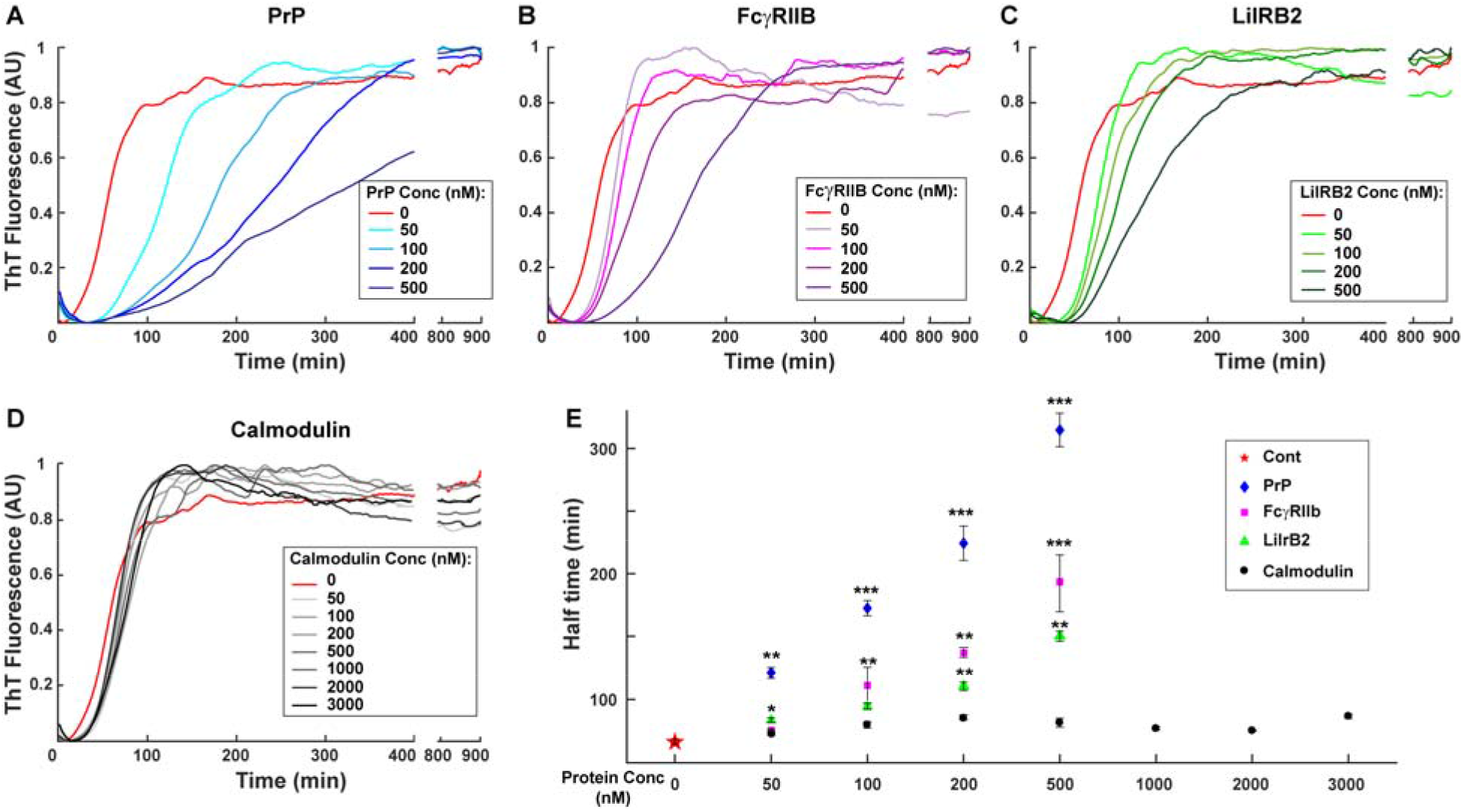
FcγRIIb and LilrB2 affect the kinetics of Aβ polymerization in a manner similar to PrP. ThT curves for polymerization of unlabeled Aβ (5 μM) in the presence of increasing concentrations of recombinant PrP **(A)**, FcγRIIb **(B)**, LilrB2 **(C)** and calmodulin **(D)**. **(E)** Effect of receptors on the half-times for Aβ polymerization, derived from the data in panels A-D. Data represent mean ± S.E. Half-time values that are significantly different from control are indicated: *P<0.05, **P<0.01, and ***P< 0.001 (Student’s t-test).

We also analyzed the effect of these receptor proteins on Aβ fibril formation by super-resolution microscopy (Fig. 7). Strikingly, we found that both FcγRIIb and LilrB2 affect Aβ fibril length and number in a manner very similar to PrP. Thus, Aβ aggregates formed in the presence of FcγRIIb and LilrB2 were shorter and more numerous than under control conditions (Fig 7G-L and M-R respectively). As is the case for PrP, these effects were concentration-dependent, and occurred with substoichiometric levels of the receptor proteins. In contrast, fibrils formed in the presence of different concentrations of calmodulin, were very similar to control fibrils in terms of their length and number (Fig. 7A-F).

**Fig. 7.**
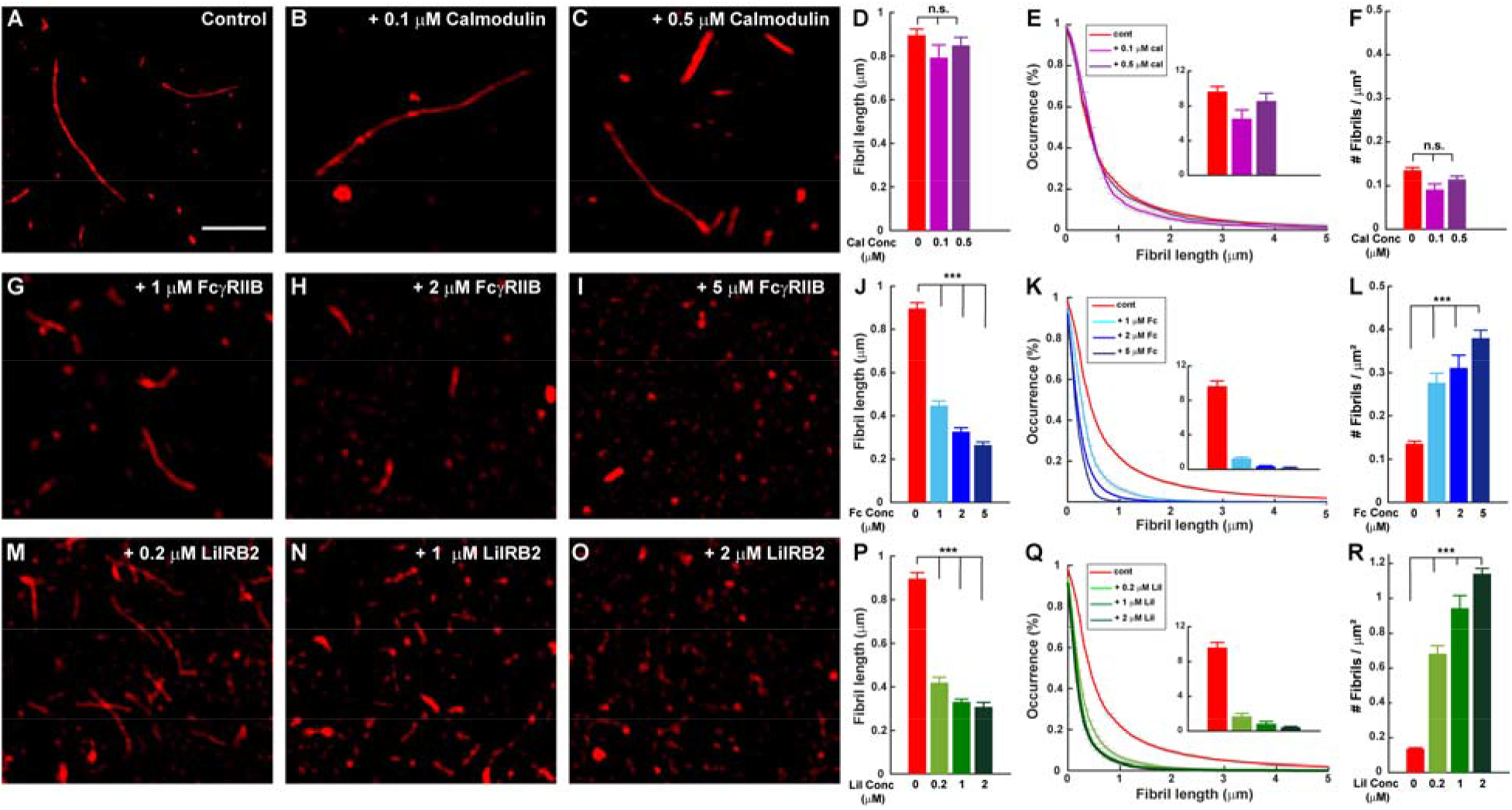
FcγRIIb and LilrB2 promote formation of shorter, more numerous Aβ fibrils. Aβ-Cy5 monomer (20 μM) was polymerized for 24 h under control conditions, or in the presence of calmodulin (0.1 and 0.5 μM), FcγRIIb (1, 2, and 5 μM), or LilrB2 (0.2, 1, and 2 μM). Panels **A-C**, **G-I**, and **M-O** show SIM images of the resulting fibrils. Scale bar in panel A is 2 μm. Panels **D**, **J**, and **P** show mean fibril length under each condition. Panels **E**, **K**, and **Q** show cumulative distributions of fibril length; the insets indicates the number of fibrils larger than 2 μm. Panels **F**, **L**, and **R**show the number of fibrils/μm^2^. Data represent mean ± S.E. ** P<0.01 and ***P< 0.001 (Student’s t-test). n.s., not statistically significant.

## DISCUSSION

There has been considerable interest in identifying the cell-surface receptors that mediate the neurotoxic effects of Aβ oligomers, in part because of the possibility that small molecules targeting these receptors, or the downstream pathways they activate, could be used as therapeutic agents to treat AD. At least 10 different cell-surface proteins have been proposed to act as Aβ receptors^8,9^. However, there has been controversy about the functional relevance of these receptors, and how much each contributes to the pathogenesis of AD. Part of the uncertainty on this subject derives from lack of detailed structural and molecular information about how these receptors interact with different kinds of Aβ aggregates. While many previous studies have characterized the binding reaction using biochemical or cell-based methods, we have employed super-resolution microscopy to directly visualize Aβ-receptor interactions at nanoscale resolution. This approach has allowed us to elucidate a unique structural mechanism by which one important receptor, PrP^C^, influences the Aβ assembly process, and how it interacts with both Aβ fibrils as well as neurotoxic Aβ oligomers. By extending our studies to two additional Aβ receptors, we have revealed common mechanistic principles that have important implications for Aβ neurotoxic signaling, as well as the development of potential therapies for AD.

In our previously published study^22^, we showed that PrP, in sub-stoichiometric amounts, potently inhibits the process of Aβ polymerization, as monitored by ThT binding and biochemical assays. Based on mathematical modeling of polymerization kinetics, we demonstrated that PrP^C^ specifically inhibits the elongation step of Aβ fibril growth, and we suggested that this might result from PrP binding to the growing ends of fibrils, thereby blocking their further elongation. However, this study did not directly measure the effect of PrP on fibril lengths or elongation rates, and did not visualize the localization of PrP on individual fibrils. In the present study, we have greatly extended this previous work using single molecule, super-resolution imaging.

Taken together, our results suggest a model (Fig. 8A) in which PrP^C^ inhibits the elongation step of Aβ fibril growth by binding specifically to only one end (the more rapidly growing end) of each fibril, thereby preventing further addition of monomers at that end. Under these conditions, fibril growth can proceed only at the slowly growing end. Multiple lines of evidence reported here support this model. First, fibrils formed in the presence of sub-stoichiometric amounts of PrP are shorter and grow more slowly than under control conditions. Second, PrP causes a significant increase in the total number of fibrils present at any given time. This latter effect is seen with other proteins that inhibit the elongation step of fibril growth, and is predicted by the kinetic models of polymerization based on an increased flux of monomers into secondary nucleation events ^34,37^. Third, seeded polymerization experiments, which allow measurement of elongation at the two ends of the fibril separately, demonstrate that PrP completely blocks elongation at one end (the normally fast-growing end), leaving elongation to proceed exclusively at the slow-growing end. Fourth, PrP is localized exclusively at one end of individual Aβ fibrils, and this end corresponds to the fast-growing end under control conditions.

**Fig. 8.**
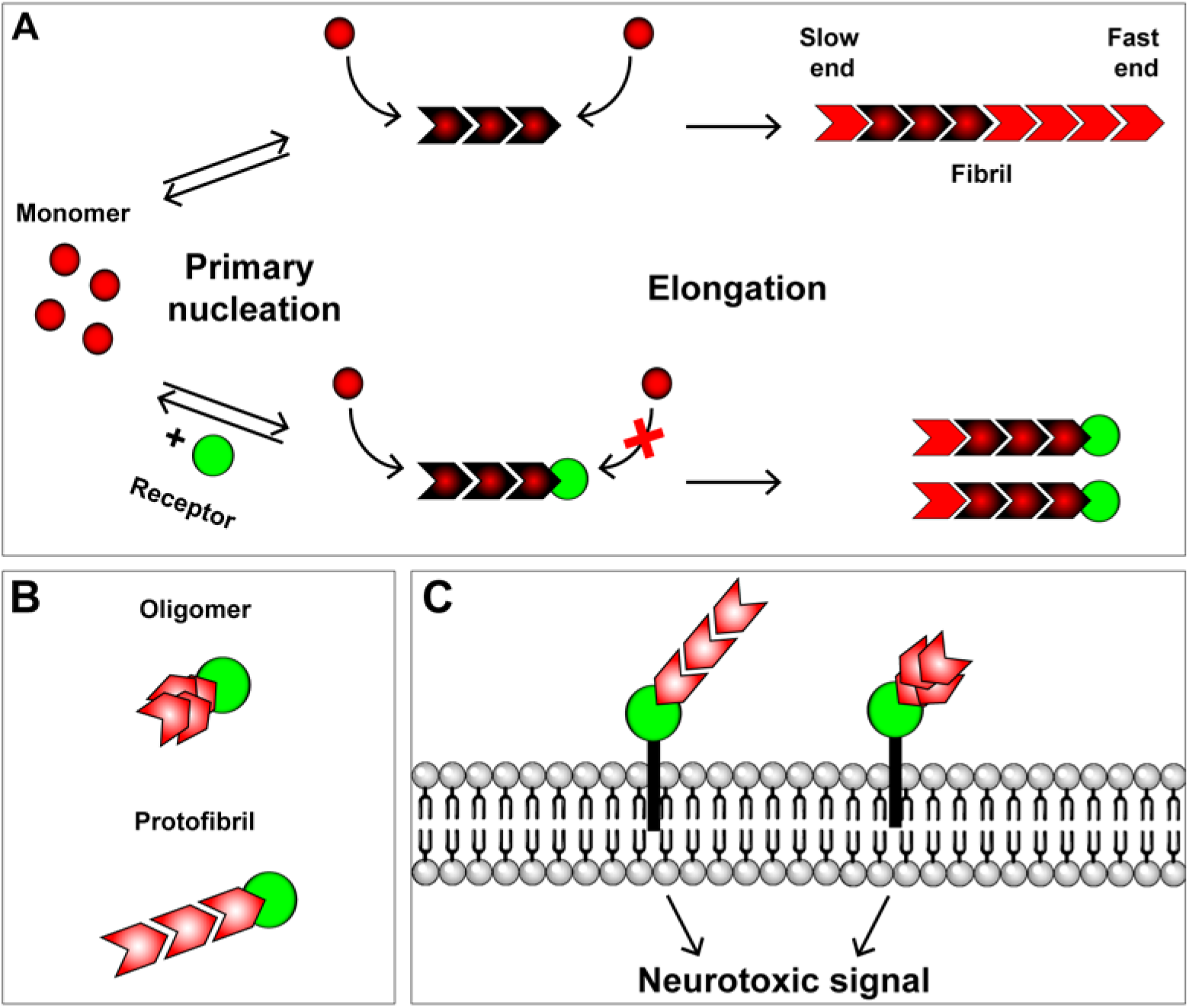
Models for the interaction of receptors with Aβ fibrils, protofibrils, and oligomers. **(A)** Schematic showing primary nucleation and elongation steps in the Aβ polymerization process in the absence (upper pathway) and the presence (lower pathway) of a receptor protein, such as PrP^C^, FcγRIIb, or LilrB2. The receptor binds to the fast-growing end of the fibril, blocking elongation at that end, and restricting elongation to the slow-growing end. Secondary nucleation events are not depicted. **(B)** Interaction of a receptor with neurotoxic oligomers and protofibrils. Receptors bind to one end/edge of these assemblies, possibly recognizing the same structural determinant present at the fast-growing end of fibrils. **(C)** Binding of Aβ oligomers and protofibrils to membrane-anchored receptors initiates neurotoxic signaling.

The model proposed here is consistent with recently a published atomic structure of Aβ(1-42) fibrils determined by cryo-EM in conjunction with solid-state NMR and X-ray crystallography^54^. This study shows that fibrils have two structurally distinct ends, termed “groove” and “ridge”, based on the binding interface presented to newly added monomers at each end. It was suggested that this structural dimorphism accounts for the polarity of Aβ fibril growth, which was recently demonstrated^43^. Another, earlier study also presented an Aβ(1-42) fibril structure that displayed a structural polarity^55^. Based on these studies, we postulate that PrP is able to bind specifically to the fast-growing end of the fibril because it recognizes a unique structural interface presented at that end. Our previous study suggested that this is a high affinity interaction with a Kd of 47.6 nM^22^. In that study, we showed that, although the flexible, N-terminal domain of PrP contains both of the identified Aβ binding sites, the structured C-terminal domain is required in order for PrP to inhibit the Aβ polymerization reaction. Thus, both domains of PrP may contribute structurally to interaction with the end of the Aβ fibril.

There is a great deal of evidence that small oligomeric forms of Aβ, rather than long amyloid fibrils, are the neurotoxic species primarily responsible for the synaptic loss and cognitive decline in AD^5–7^. We therefore sought to analyze the interaction of PrP with two oligomeric forms of Aβ, ADDLs and protofibrils, both of which we have confirmed are highly neurotoxic in a dendritic spine retraction assay. Using super-resolution imaging, we localized PrP to one end of short protofibrils, similar to its localization on longer, mature fibrils. On ADDLs, which are globular or ellipsoid in shape without clearly identifiable ends, it was more difficult to resolve the precise location of PrP binding, particularly since these assemblies are at the resolution limit of dSTORM imaging. Nevertheless, we found a statistically significant tendency for PrP to localize eccentrically, toward one edge of the ADDL aggregate. A similar eccentric localization of PrP was observed on small oligomers of Aβ that were normally present during the course of the Aβ polymerization reaction, particularly in the presence of PrP. We postulate that the eccentric localization of PrP on ADDLs and small oligomers may reflect an intrinsic asymmetry in their structure. This suggestion is consistent with the recently published cryo-EM structure of Aβ fibrils^54^, which indicated that the “groove” and “ridge” ends are formed once the fibril reaches a minimum size of six subunits. These considerations raise the interesting possibility that the structural determinants recognized by PrP on neurotoxic oligomers of Aβ may be similar to the ones PrP recognizes on the fast-growing end of polymerizing fibrils (Fig. 8B). In this case, the potent neurotoxicity of oligomers would result from the fact that oligomers, in contrast to fibrils, present a high molar concentration of “end-specific” structural determinants to PrP or other cell-surface receptors (Fig. 8C).

Most previous literature on the interaction of PrP^C^ and Aβ has focused on oligomeric forms of Aβ, particularly synthetic ADDLs, since these are the forms thought to be most relevant to neurotoxicity *in vivo*^10,19–22^. Only a few studies have analyzed the interaction of PrP with Aβ fibrils, or its effect on polymerization or assembly processes. Several different effects of PrP on Aβ aggregation have been described, including inhibition of fibril formation^56^, bundling^57^ or fragmentation^58,59^ of fibrils, and trapping of Aβ in an oligomeric state within PrP-containing complexes^58–60^. In a particularly relevant example, it was reported that protofibrils (like those used here) were the most neurotoxic forms of Aβ in an LTP suppression assay, and were the forms that bound most avidly to PrP^21^. However, none of these previous studies localized PrP to particular sites on Aβ assemblies, or identified specific steps in the assembly process that were affected.

Previous studies have identified several other proteins that act as chaperones affecting particular steps in the polymerization pathway of Aβ or other amyloidogenic proteins^34^. For example, Ssa1, an Hsp70-type chaperone in yeast, has been shown to block elongation of fibrils formed by the prion-like protein, Ure2p^61^. Like other biological polymers^62^, amyloid fibrils have been shown to elongate in a polarized fashion, with the two ends growing at different rates, reflecting distinct structural interfaces at each end^43–45,63^. PrP is, to our knowledge, the first factor to be identified that acts as an end-specific inhibitor of amyloid fibril elongation.

The experiments presented here suggest that two other putative Aβ receptors, FcγRIIb and LilrB2, interact with Aβ via a mechanism similar to that of PrP, involving selective inhibition of elongation. These receptors, which are expressed in both immune cells and neurons, have been shown to bind Aβ oligomers, and their genetic ablation reduces oligomer-induced synaptic toxicity in hippocampal slices^11,12,51^. We found that the extracellular domains of both FcγRIIb and LilrB2, which comprise the known Aβ binding sites, inhibit Aβ polymerization similarly to PrP: in sub-stoichiometric amounts, they increased the polymerization half-time as measured by ThT fluorescence, and they created shorter and more numerous fibrils as monitored by super-resolution imaging. These data suggest a common molecular mechanism by which different receptors interact with Aβ fibrils, and perhaps with neurotoxic oligomers as well. PrP^C^ has been implicated as a cell surface receptor for other toxic oligomers, including those composed of tau and α-synuclein^64–66^, and it will be of interest to determine whether PrP^C^ interacts with these proteins and affects their fibrilization by mechanisms similar to those we have shown here for Aβ.

Interestingly, available evidence suggests that the signal transduction pathways stimulated by Aβ oligomer binding to PrP^C^, FcγRIIb, and LilrB2 may all be different^12,25,26,51^. Thus, binding of multiple receptors to a common set of structural determinants on Aβ oligomers may activate an array of different signaling mechanisms, which mediate distinct aspects of the synaptotoxic response. PrP^C^ also serves as a cell-surface receptor mediating the synaptotoxic activity of PrP^Sc^, the infectious form of PrP. However, our evidence indicates that the signaling pathways initiated by PrP^Sc^ and Aβ may not be identical^50^.

Our results raise the previously unappreciated possibility that PrP^C^, as well as other cell surface receptors, in addition to serving as signal transduction elements, could directly influence the Aβ assembly process, favoring the formation of smaller, more numerous fibrils or protofibrils with enhanced neurotoxicity. Several endogenous proteins and non-protein molecules have been found to influence the formation and stability of Aβ aggregates, thereby modulating their neurotoxicity^37,67^. PrP^C^ in particular is an abundant brain protein, which is likely present in sufficient amounts in relation to Aβ to influence the assembly of Aβ aggregates, particularly since it acts sub-stoichiometrically. The fact that PrP may promote formation of smaller, more neurotoxic aggregates would argue against the proposed strategy of using soluble, recombinant PrP as a drug to treat AD^68^.

Current therapies for AD are focused primarily on lowering levels of Aβ, either by inhibiting its synthesis or enhancing its degradation^69^. These therapies have met with little success in recent clinical trials. The results presented here raise the possibility of a novel therapeutic approach based on blocking interactions of neurotoxic forms of Aβ with its cellular receptors, thereby inhibiting downstream signaling pathways engaged by these receptors. Our work suggests that this approach may be feasible, based our conclusion that there is a specific structural interface on Aβ fibril ends, as well as on neurotoxic protofibrils and oligomers, which is recognized by multiple Aβ receptors. By defining this interface at the atomic level, it may be possible to design small molecules that specifically block receptor-Aβ interactions or signaling mechanisms, thereby preventing AD pathology. This structural information may also inform creation of diagnostic reagents specific for neurotoxic forms of Aβ.

## ACKNOWLEDGMENTS

This work was supported by NIH grant R01 NS065244 to D.A.H.; and an Alzheimer’s Association Research Fellowship (2018-AARF-589708) to L.A.. Super-resolution microscopy was performed at the Harvard Center for Biological Imaging.

## AUTHOR CONTRIBUTION

D.A.H. and L.A. conceived the project. L.A. performed the experiments. L.A. and D.A.H. designed the experiments, analyzed the data and wrote the paper.

## CONFILICT OF INTEREST

The authors declare that they have no conflicts of interest with the contents of this article.

## REFERENCES

1 Jack, C. R., Jr. et al. NIA-AA Research Framework: Toward a biological definition of Alzheimer’s disease. Alzheimer’s & dementia: the journal of the Alzheimer’s Association 14, 535–562, doi:10.1016/j.jalz.2018.02.018 (2018).

2 Long, J. M. & Holtzman, D. M. Alzheimer Disease: An Update on Pathobiology and Treatment Strategies. Cell 179, 312–339, doi:10.1016/j.cell.2019.09.001 (2019).

3 Selkoe, D. J. Alzheimer’s disease. Cold Spring Harbor perspectives in biology 3, doi:10.1101/cshperspect.a004457 (2011).

4 Serrano-Pozo, A., Frosch, M. P., Masliah, E. & Hyman, B. T. Neuropathological alterations in Alzheimer disease. Cold Spring Harbor perspectives in medicine 1, a006189, doi:10.1101/cshperspect.a006189 (2011).

5 Shankar, G. M. et al. Amyloid-beta protein dimers isolated directly from Alzheimer’s brains impair synaptic plasticity and memory. Nature medicine 14, 837–842, doi:10.1038/nm1782 (2008).

6 Walsh, D. M. & Selkoe, D. J. A beta oligomers - a decade of discovery. Journal of neurochemistry 101, 1172–1184, doi:10.1111/j.1471-4159.2006.04426.x (2007).

7 Lacor, P. N. et al. Aβ oligomer-induced aberrations in synapse composition, shape, and density provide a molecular basis for loss of connectivity in Alzheimer’s disease. J. Neurosci. 27, 796–807 (2007).

8 Smith, L. M. & Strittmatter, S. M. Binding Sites for Amyloid-beta Oligomers and Synaptic Toxicity. Cold Spring Harbor perspectives in medicine 7, doi:10.1101/cshperspect.a024075 (2017).

9 Smith, L. M., Kostylev, M. A., Lee, S. & Strittmatter, S. M. Systematic and standardized comparison of reported amyloid-β receptors for sufficiency, affinity, and Alzheimer’s disease relevance. J. Biol. Chem. 294, 6042–6053, doi:10.1074/jbc.RA118.006252 (2019).

10 Lauren, J., Gimbel, D. A., Nygaard, H. B., Gilbert, J. W. & Strittmatter, S. M. Cellular prion protein mediates impairment of synaptic plasticity by amyloid-beta oligomers. Nature 457, 1128–1132, doi:10.1038/nature07761 (2009).

11 Kam, T. I. et al. FcgammaRIIb mediates amyloid-beta neurotoxicity and memory impairment in Alzheimer’s disease. The Journal of clinical investigation 123, 2791–2802, doi:10.1172/jci66827 (2013).

12 Kim, T. et al. Human LilrB2 is a beta-amyloid receptor and its murine homolog PirB regulates synaptic plasticity in an Alzheimer’s model. Science (New York, N.Y.) 341, 1399–1404, doi:10.1126/science.1242077 (2013).

13 Cisse, M. et al. Reversing EphB2 depletion rescues cognitive functions in Alzheimer model. Nature 469, 47–52, doi:10.1038/nature09635 (2011).

14 Zhao, Y. et al. Amyloid Beta Peptides Block New Synapse Assembly by Nogo Receptor-Mediated Inhibition of T-Type Calcium Channels. Neuron 96, 355–372.e356, doi:10.1016/j.neuron.2017.09.041 (2017).

15 Cisse, M. et al. Ablation of cellular prion protein does not ameliorate abnormal neural network activity or cognitive dysfunction in the J20 line of human amyloid precursor protein transgenic mice. J. Neurosci. 31, 10427–10431, doi:31/29/10427 [pii]10.1523/JNEUROSCI.1459-11.2011 (2011).

16 Balducci, C. et al. Synthetic amyloid-beta oligomers impair long-term memory independently of cellular prion protein. Proceedings of the National Academy of Sciences of the United States of America 107, 2295–2300, doi:10.1073/pnas.0911829107 (2010).

17 Calella, A. M. et al. Prion protein and Abeta-related synaptic toxicity impairment. EMBO molecular medicine 2, 306–314, doi:10.1002/emmm.201000082 (2010).

18 Kessels, H. W., Nguyen, L. N., Nabavi, S. & Malinow, R. The prion protein as a receptor for amyloid-beta. Nature 466, E3–4; discussion E4-5, doi:10.1038/nature09217 (2010).

19 Chen, S., Yadav, S. P. & Surewicz, W. K. Interaction between human prion protein and amyloid-beta (Abeta) oligomers: role OF N-terminal residues. The Journal of biological chemistry 285, 26377–26383, doi:10.1074/jbc.M110.145516 (2010).

20 Fluharty, B. R. et al. An N-terminal fragment of the prion protein binds to amyloid-beta oligomers and inhibits their neurotoxicity in vivo. The Journal of biological chemistry 288, 7857–7866, doi:10.1074/jbc.M112.423954 (2013).

21 Nicoll, A. J. et al. Amyloid-beta nanotubes are associated with prion protein-dependent synaptotoxicity. Nature communications 4, 2416, doi:10.1038/ncomms3416 (2013).

22 Bove-Fenderson, E., Urano, R., Straub, J. E. & Harris, D. A. Cellular prion protein targets amyloid-beta fibril ends via its C-terminal domain to prevent elongation. The Journal of biological chemistry 292, 16858–16871, doi:10.1074/jbc.M117.789990 (2017).

23 Ganzinger, K. A. et al. Single-molecule imaging reveals that small amyloid-beta1-42 oligomers interact with the cellular prion protein (PrP(C)). Chembiochem 15, 2515–2521, doi:10.1002/cbic.201402377 (2014).

24 Freir, D. B. et al. Interaction between prion protein and toxic amyloid β assemblies can be therapeutically targeted at multiple sites. Nat. Commun. 2, 336, doi:10.1038/ncomms1341ncomms1341 [pii] (2011).

25 Um, J. W. et al. Alzheimer amyloid-beta oligomer bound to postsynaptic prion protein activates Fyn to impair neurons. Nature neuroscience 15, 1227–1235, doi:10.1038/nn.3178 (2012).

26 Um, J. W. et al. Metabotropic glutamate receptor 5 is a coreceptor for Alzheimer abeta oligomer bound to cellular prion protein. Neuron 79, 887–902, doi:10.1016/j.neuron.2013.06.036 (2013).

27 Haas, L. T. & Strittmatter, S. M. Oligomers of Amyloid beta Prevent Physiological Activation of the Cellular Prion Protein-Metabotropic Glutamate Receptor 5 Complex by Glutamate in Alzheimer Disease. The Journal of biological chemistry 291, 17112–17121, doi:10.1074/jbc.M116.720664 (2016).

28 Haas, L. T., Kostylev, M. A. & Strittmatter, S. M. Therapeutic molecules and endogenous ligands regulate the interaction between brain cellular prion protein (PrPC) and metabotropic glutamate receptor 5 (mGluR5). The Journal of biological chemistry 289, 28460–28477, doi:10.1074/jbc.M114.584342 (2014).

29 Gimbel, D. A. et al. Memory impairment in transgenic Alzheimer mice requires cellular prion protein. The Journal of neuroscience: the official journal of the Society for Neuroscience 30, 6367–6374, doi:10.1523/jneurosci.0395-10.2010 (2010).

30 Gunther, E. C. et al. Rescue of transgenic Alzheimer’s pathophysiology by polymeric cellular prion protein antagonists. Cell Rep. 26, 145–158 e148, doi:10.1016/j.celrep.2018.12.021 (2019).

31 Chung, E. et al. Anti-PrP^C^ monoclonal antibody infusion as a novel treatment for cognitive deficits in an Alzheimer’s disease model mouse. BMC Neurosci. 11, 130, doi:1471-2202-11-130 [pii] 10.1186/1471-2202-11-130 (2010).

32 van Dyck, C. H. et al. Effect of AZD0530 on Cerebral Metabolic Decline in Alzheimer Disease: A Randomized Clinical Trial. JAMA Neurol, doi:10.1001/jamaneurol.2019.2050 (2019).

33 Cohen, S. I. et al. Proliferation of amyloid-beta42 aggregates occurs through a secondary nucleation mechanism. Proceedings of the National Academy of Sciences of the United States of America 110, 9758–9763, doi:10.1073/pnas.1218402110 (2013).

34 Arosio, P. et al. Kinetic analysis reveals the diversity of microscopic mechanisms through which molecular chaperones suppress amyloid formation. Nature communications 7, 10948, doi:10.1038/ncomms10948 (2016).

35 Cohen, S. I., Vendruscolo, M., Dobson, C. M. & Knowles, T. P. From macroscopic measurements to microscopic mechanisms of protein aggregation. Journal of molecular biology 421, 160–171, doi:10.1016/j.jmb.2012.02.031 (2012).

36 Beeg, M. et al. Clusterin Binds to Abeta1-42 Oligomers with High Affinity and Interferes with Peptide Aggregation by Inhibiting Primary and Secondary Nucleation. The Journal of biological chemistry 291, 6958–6966, doi:10.1074/jbc.M115.689539 (2016).

37 Cohen, S. I. A. et al. A molecular chaperone breaks the catalytic cycle that generates toxic Abeta oligomers. Nature structural & molecular biology 22, 207–213, doi:10.1038/nsmb.2971 (2015).

38 Habchi, J. et al. Systematic development of small molecules to inhibit specific microscopic steps of Abeta42 aggregation in Alzheimer’s disease. Proceedings of the National Academy of Sciences of the United States of America 114, E200–E208, doi:10.1073/pnas.1615613114 (2017).

39 Munke, A. et al. Phage display and kinetic selection of antibodies that specifically inhibit amyloid self-replication. Proceedings of the National Academy of Sciences of the United States of America 114, 6444–6449, doi:10.1073/pnas.1700407114 (2017).

40 Rust, M. J., Bates, M. & Zhuang, X. Sub-diffraction-limit imaging by stochastic optical reconstruction microscopy (STORM). Nature methods 3, 793–795, doi:10.1038/nmeth929 (2006).

41 Kner, P., Chhun, B. B., Griffis, E. R., Winoto, L. & Gustafsson, M. G. Super-resolution video microscopy of live cells by structured illumination. Nature methods 6, 339–342, doi:10.1038/nmeth.1324 (2009).

42 Hellstrand, E., Boland, B., Walsh, D. M. & Linse, S. Amyloid beta-protein aggregation produces highly reproducible kinetic data and occurs by a two-phase process. ACS chemical neuroscience 1, 13–18, doi:10.1021/cn900015v (2010).

43 Young, L. J., Kaminski Schierle, G. S. & Kaminski, C. F. Imaging Abeta(1-42) fibril elongation reveals strongly polarised growth and growth incompetent states. Physical chemistry chemical physics: PCCP 19, 27987–27996, doi:10.1039/c7cp03412a (2017).

44 Pinotsi, D. et al. Direct observation of heterogeneous amyloid fibril growth kinetics via two-color super-resolution microscopy. Nano letters 14, 339–345, doi:10.1021/nl4041093 (2014).

45 DePace, A. H. & Weissman, J. S. Origins and kinetic consequences of diversity in Sup35 yeast prion fibers. Nature structural biology 9, 389–396, doi:10.1038/nsb786 (2002).

46 Ban, T. et al. Direct observation of Abeta amyloid fibril growth and inhibition. Journal of molecular biology 344, 757–767, doi:10.1016/j.jmb.2004.09.078 (2004).

47 Lambert, M. P. et al. Diffusible, nonfibrillar ligands derived from Abeta1-42 are potent central nervous system neurotoxins. Proceedings of the National Academy of Sciences of the United States of America 95, 6448–6453 (1998).

48 Caughey, B. & Lansbury, P. T. Protofibrils, pores, fibrils, and neurodegeneration: separating the responsible protein aggregates from the innocent bystanders. Annu. Rev. Neurosci. 26, 267–298 (2003).

49 Fang, C., Imberdis, T., Garza, M. C., Wille, H. & Harris, D. A. A Neuronal Culture System to Detect Prion Synaptotoxicity. PLoS pathogens 12, e1005623, doi:10.1371/journal.ppat.1005623 (2016).

50 Fang, C. et al. Prions activate a p38 MAPK synaptotoxic signaling pathway. PLoS Pathog. 14, e1007283, doi:10.1371/journal.ppat.1007283 (2018).

51 Kam, T. I. et al. FcgammaRIIb-SHIP2 axis links Abeta to tau pathology by disrupting phosphoinositide metabolism in Alzheimer’s disease model. eLife 5, doi:10.7554/eLife.18691 (2016).

52 Pritchard, N. R. & Smith, K. G. B cell inhibitory receptors and autoimmunity. Immunology 108, 263–273 (2003).

53 Syken, J., Grandpre, T., Kanold, P. O. & Shatz, C. J. PirB restricts ocular-dominance plasticity in visual cortex. Science (New York, N.Y.) 313, 1795–1800, doi:10.1126/science.1128232 (2006).

54 Gremer, L. et al. Fibril structure of amyloid-beta(1-42) by cryo-electron microscopy. Science (New York, N.Y.) 358, 116–119, doi:10.1126/science.aao2825 (2017).

55 Luhrs, T. et al. 3D structure of Alzheimer’s amyloid-β(1-42) fibrils. Proc. Natl. Acad. Sci. USA 102, 17342–17347, doi:10.1073/pnas.0506723102 (2005).

56 Nieznanski, K., Choi, J. K., Chen, S., Surewicz, K. & Surewicz, W. K. Soluble prion protein inhibits amyloid β (Aβ) fibrillization and toxicity. J. Biol. Chem. 287, 33104–33108, doi:C112.400614 [pii] 10.1074/jbc.C112.400614 (2012).

57 Nieznanski, K., Surewicz, K., Chen, S., Nieznanska, H. & Surewicz, W. K. Interaction between prion protein and Abeta amyloid fibrils revisited. ACS chemical neuroscience 5, 340–345, doi:10.1021/cn500019c (2014).

58 Younan, N. D., Sarell, C. J., Davies, P., Brown, D. R. & Viles, J. H. The cellular prion protein traps Alzheimer’s Aβ in an oligomeric form and disassembles amyloid fibers. FASEB J. 27, 1847–1858, doi:10.1096/fj.12-222588fj.12-222588 [pii] (2013).

59 Younan, N. D., Chen, K. F., Rose, R. S., Crowther, D. C. & Viles, J. H. Prion protein stabilizes amyloid-beta (Abeta) oligomers and enhances Abeta neurotoxicity in a Drosophila model of Alzheimer’s disease. The Journal of biological chemistry 293, 13090–13099, doi:10.1074/jbc.RA118.003319 (2018).

60 Rosener, N. S. et al. A d-enantiomeric peptide interferes with heteroassociation of amyloid-beta oligomers and prion protein. The Journal of biological chemistry 293, 15748–15764, doi:10.1074/jbc.RA118.003116 (2018).

61 Xu, L. Q. et al. Influence of specific HSP70 domains on fibril formation of the yeast prion protein Ure2. Philos. Trans. R. Soc. Lond. B Biol. Sci. 368, 20110410, doi:10.1098/rstb.2011.0410 (2013).

62 Oosawa, F. & Kasai, M. A theory of linear and helical aggregations of macromolecules. Journal of molecular biology 4, 10–21, doi:10.1016/s0022-28366280112-0 (1962).

63 Ban, T., Hamada, D., Hasegawa, K., Naiki, H. & Goto, Y. Direct observation of amyloid fibril growth monitored by thioflavin T fluorescence. The Journal of biological chemistry 278, 16462–16465, doi:10.1074/jbc.C300049200 (2003).

64 Ondrejcak, T. et al. Cellular Prion Protein Mediates the Disruption of Hippocampal Synaptic Plasticity by Soluble Tau In Vivo. The Journal of neuroscience: the official journal of the Society for Neuroscience 38, 10595–10606, doi:10.1523/JNEUROSCI.1700-18.2018 (2018).

65 Ferreira, D. G. et al. α-Synuclein interacts with PrP^C^ to induce cognitive impairment through mGluR5 and NMDAR2B. Nat. Neurosci. 20, 1569–1579, doi:10.1038/nn.4648 (2017).

66 Aulic, S. et al. α-Synuclein amyloids hijack prion protein to gain cell entry, facilitate cell-to-cell spreading and block prion replication. Sci. Rep. 7, 10050, doi:10.1038/s41598-017-10236-x (2017).

67 Liu, C. C. et al. Neuronal heparan sulfates promote amyloid pathology by modulating brain amyloid-beta clearance and aggregation in Alzheimer’s disease. Sci Transl Med 8, 332ra344, doi:10.1126/scitranslmed.aad3650 (2016).

68 Scott-McKean, J. J. et al. Soluble prion protein and its N-terminal fragment prevent impairment of synaptic plasticity by Abeta oligomers: Implications for novel therapeutic strategy in Alzheimer’s disease. Neurobiol Dis 91, 124–131, doi:10.1016/j.nbd.2016.03.001 (2016).

69 Golde, T. E., DeKosky, S. T. & Galasko, D. Alzheimer’s disease: The right drug, the right time. Science (New York, N.Y.) 362, 1250–1251, doi:10.1126/science.aau0437 (2018).

