## Supplementary information for "Aβ receptors specifically recognize molecular features displayed by fibril ends and neurotoxic oligomers"

### METHODS

#### Preparation of Aβ monomers

Lyophilized Aβ (1-42), Aβ-Cy5 (1-42) and Aβ-Cy3 (1-42), were synthesized by ERI Amyloid Laboratory, LLC (Oxford, CT, USA). Details of monomer perpetration can be found in Bove-Fenderson, et al. (2017)[1]. Unlabeled monomers were solubilized in 15 mM NaOH, and were then isolated by size exclusion chromatography on a Superdex 75 10/300 GL (GE Healthcare) column using PBS as the running buffer. Fractions were collected and were immediately used in ThT assays. Cy3- and Cy5-labeled peptide was solubilized in 15 mM NaOH and was used directly for ThT assays and super-resolution microscopy. The concentration of Aβ was estimated with a NanoDrop UV-visible spectrometer (Thermo Scientific) by reading the sample absorbance at 214 nm and applying Beer's Law with an extinction coefficient of 76848 M-1cm- 1.

#### Preparation of ADDLs and protofibrils

Fluorescently labelled ADDLs were prepared using a standard protocol[2,3] in which lyophilized Aβ peptide was solubilized in HFIP and then dried to a film. The ratio between labeled and unlabeled peptide was 1:10. The film was then solubilized in DMSO before dilution to a concentration of 100 µM in phenol red-free Ham's F12 medium (DMSO 2% v/v), followed by incubation at room temperature for 16 hrs. To prepare protofibrils[4], ADDL samples were incubated at room temperature for either 3 days (protofibril preparation 1) or 7 days (protofibril preparation 2).

#### Recombinant PrP

Full-length mouse PrP (23-230) was produced and purified as described previously[1]. *E. coli* strain BL21 Star was transformed with the pJ411 vector expressing murine PrP23-230. Cells were lysed, and then PrP was purified with an ÄKTA purification system (GE Healthcare) using a $Ni^{2+}$-immobilized metal ion affinity column. Protein was eluted from the $Ni^{2+}$ immobilized metal ion affinity column with 5 M guanidine HCl, 0.1 M Tris acetate, 0.1 M potassium phosphate (pH 4.5) while monitoring $A_{280}$. Fractions spanning the elution peak were combined, and the pH was raised to 8 by titration with potassium acetate. The pooled samples were then desalted into 20 mM potassium acetate, pH 5.5 using a HiPrep 26/10 desalting column (GE Healthcare), and PrP was purified by reverse-phase HPLC using a C4 column (Grace/Vydac). Fractions containing the purified protein were pooled, lyophilized, and stored at −80 °C for future use.

For fluorescent labeling, PrP was prepared with a cysteine residue substituted for a glycine residue at position 34 (G34C). Lyophilized PrP G34C was dissolved in 20 mM potassium acetate, pH 5.5 to a concentration of 100 µM. Alexa Fluor 555 or 488 C2 maleimide (ThermoFisher Scientific) was added dropwise with stirring from a stock solution of 1 mM in water, to a final ratio of 1:10 (protein:dye). This solution was incubated at room temperature for two hours. One ml of the solution was then injected into an analytical C3 column (Zorbax 300SB C3, Agilent) on an Agilent

1200 Infinity HPLC system, and the peptide peak/dye was collected and lyophilized. Confirmation of successful linkage was made by MALDI-TOF mass spectrometry.

**Other recombinant Proteins**

Recombinant FcγRIIb and LilrB2 (extracellular domains) were purchased from Novoprotein (C444) and R&D systems (8429-T4), respectively.

**ThT assay for Aβ polymerization**

Kinetic assays for Aβ polymerization were conducted as described previously[1,5,6]. Aβ monomers were diluted to a concentration of 5 to 20 μM in PBS, and 10 μM ThT was added. Recombinant proteins were added from a 1 mg/ml stock in water at the indicated concentrations. To follow ThT binding, 100 μl samples were placed in 96-well, half-volume, low-binding plates (Corning 3881), and florescence was read in a Synergy H1 Multi-Mode Microplate Reader (BioTek) every 2 min at 37°C (excitation 440 nm, emission 480 nm).

**Preparation of Aβ samples for super-resolution microscopy**

Fluorescently labelled Aβ fibrils were formed by polymerizing 100% Cy5-labeled Aβ 1-42 monomers. Labeled monomers were diluted to a concentration of 10 to 20 μM in PBS, followed by incubation at 37°C for 24 h to 1 week. Where indicated, recombinant proteins were added to monomeric solution at the starting point of the polymerization reaction.

For seeding assays, fibrils prepared from Aβ-Cy5, as described above, were diluted in PBS to a monomer-equivalent concentration of 20 μM, and were sheared by sonication (30 sec on a 50% duty cycle, Branson 1800) to yield seeds. To initiate seeded growth, freshly prepared Aβ-Cy3 monomer (20 μM) was added to an equal volume of the seed solution, and incubated for 24 h at 37°C.

For three-color imaging, fibrils were first immobilized on antibody-coated wells. Glass-botom, multi-well plates (Lab-Tek) were sequentially cleaned in 1M HCl, 70% ethanol, and 1M KOH. After extensive rinsing with ultrapure water, plates were dried with $N_2$. The plates were then treated with an antibody against amyloid-β (6E10, mouse monoclonal primary, BioLegend) overnight. Seeded fibrils, prepared as described above, were incubated with PrP-AF488 for 30 min at 37°C, and were then added to the antibody-coated wells to allow the fibrils to adhere. After 30 min of incubation, wells were washed with ultrapure water to remove unbound PrP.

**Super-resolution imaging (SIM and dSTORM)**

Super-resolution microscopy was performed at the Harvard Center for Biological Imaging (HCBI) (Cambridge, MA, USA) using a Zeiss ELYRA microscope, which is capable of performing both dSTORM and SIM imaging. This microscope is equipped with 488 nm, 561 nm, and 638 nm laser lines, and a 100× oil immersion objective lens (NA 1.4). For SIM imaging Aβ samples were dried onto pre-washed coverslips and covered in mounting medium (Vectashield H1000, Vector laboratories). For dSTORM imaging, samples were dried onto glass-bottom multi-well plates (Lab-Tek), and were then covered in photoswitching buffer solution immediately before imaging; this solution consisted of 100 mM mercaptoethylamine (MEA) in phosphate buffered saline (PBS, pH 7.4), together with a glucose-enzyme oxygen scavenger (40 mg/ml glucose, 50 mg/ml glucose oxidase, 1 mg/ml catalase). The chamber was filled to the top, and was closed with a cap to minimize entrance of oxygen. Collected images were then processed using ZEN 2.3 software.

**Image processing**

We developed a MATLAB code to quantitate the size and number of Aβ aggregates (fibrils, protofibrils, and ADDLs) in super-resolution images. In the first step, grayscale SIM images imported to MATLAB and were smoothed by using 2×2 unit square kernel. The smoothing process helps to reduce noise within an image. The smoothed images were then binarized and converted into black-and-white image by using a threshold greater than 2× the standard deviation of the pixel value distributions. The binarization process generates sharp boundaries for each object, which were then detected by using the 'boundary' function of MATLAB. The length of each aggregate was defined by determining the maximum distance between pairs of points on the detected boundary, with these points being used to define the two ends of the aggregate. We calculated the cumulative distribution of aggregate lengths, the mean length for the distribution, and the mean number of aggregates/$\mu m^2$ in each SIM image, and took the mean values from multiple images to arrive at the final values reported in the figures.

In the case of two-color SIM images, PrP dimensions and positions were determined as described above for Aβ aggregates. The colocalization between PrP and Aβ was then computed by counting the number of PrP pixels that overlaped with Aβ aggregate pixels. This number was then normalized to total number of PrP pixels in the SIM image.

In order to quantify the localization of PrP with respect to Aβ fibril ends, we measured the minimum distance ($D_{min}$) between each fluorescent PrP spot and the closest end of the associated fibril. This distance was then normalized to the total length of the fibril ($D_{min}/L_{total}$). Random distributions were generated by using unrelated image of Aβ and PrP from two different experiments.

**Synaptotoxicity assay**

All procedures involving animals were conducted according to the United States Department of Agriculture Animal Welfare Act and the National Institutes of Health Policy on Humane Care and Use of Laboratory Animals. Hippocampal neurons were cultured from P0 pups as described[7,8]. Neurons were seeded on poly-L-lysine-coated coverslips, and after 24 h the coverslips were inverted onto an astrocyte feeder layer and maintained in NB/B27 medium until used. The astrocyte feeder layer was generated using P0 cerebral cortex. Neurons were kept in culture for 18–21 days prior to Aβ treatment.

Neurons were treated for 24 hrs with vehicle, or with 500 mM (monomer equivalent) of ADDLs, protofibrils, fibrils, followed by fixation in 4% paraformaldehyde and staining with Alexa 488-phalloidin (ThermoFischer Scientific, Waltham, MA) to visualize dendritic spines. Images were acquired using a Zeiss 880 confocal microscope with a 63x objective (N.A. = 1.4). The number dendritic spines per µm of dendrite length was determined using ImageJ software, as described previously[8].

**Statistical analysis**

Data are shown as mean ± S.E. Statistical significance of the differences between mean values was evaluated using the Student's t-test. For cases in which there were multiple comparisons, we used the Holm–Bonferroni correction to adjust the P-values. An adjusted P-value of $<0.05$ was considered to be statistically significant.

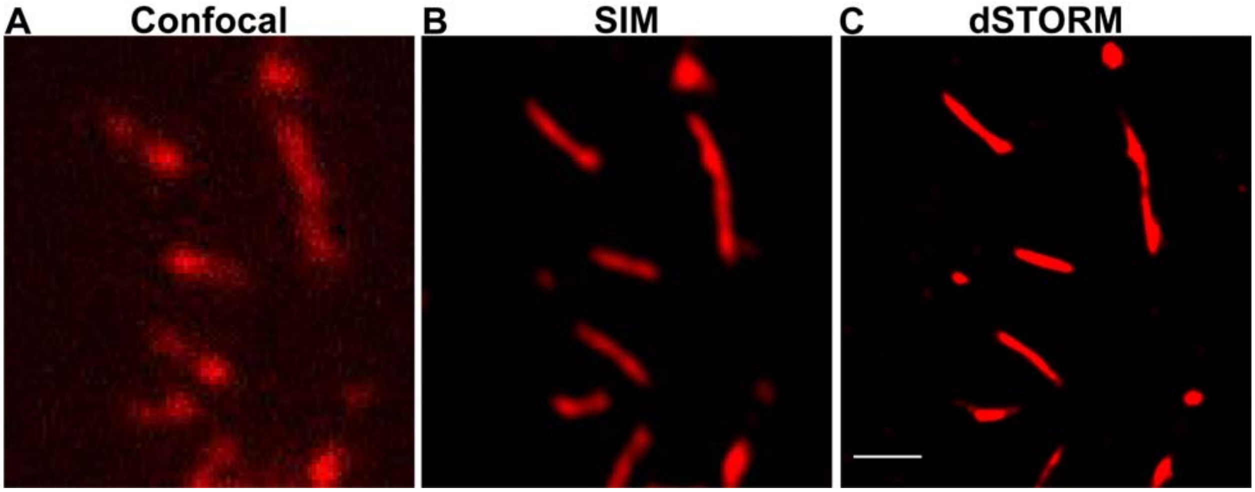


**Supplementary Fig. S1. Comparison of confocal and super-resolution imaging of Aβ fibrils.** Images of fluorescent Aβ-Cy5 fibrils acquired by laser-scanning, confocal fluorescence microscopy **(A)**, SIM **(B)**, and dSTORM **(C)**. Scale bar in panel C is 1 μm.

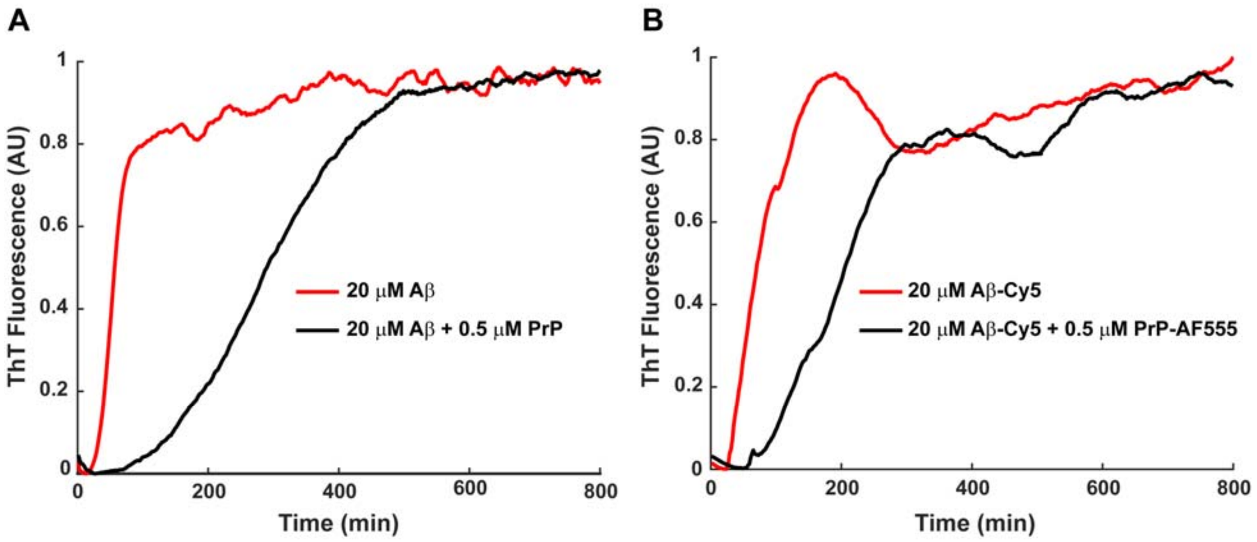


**Supplementary Fig. S2. Dye labeling does not affect the polymerization of Aβ or the inhibitory effect of PrP. (A)** ThT curves for polymerization of unlabeled Aβ (20 μM) in the presence of 0.5 μM PrP. **(B)** ThT curves for polymerization of 20 μM Aβ-Cy5 in the presence of 0.5 μM PrP-AF555.

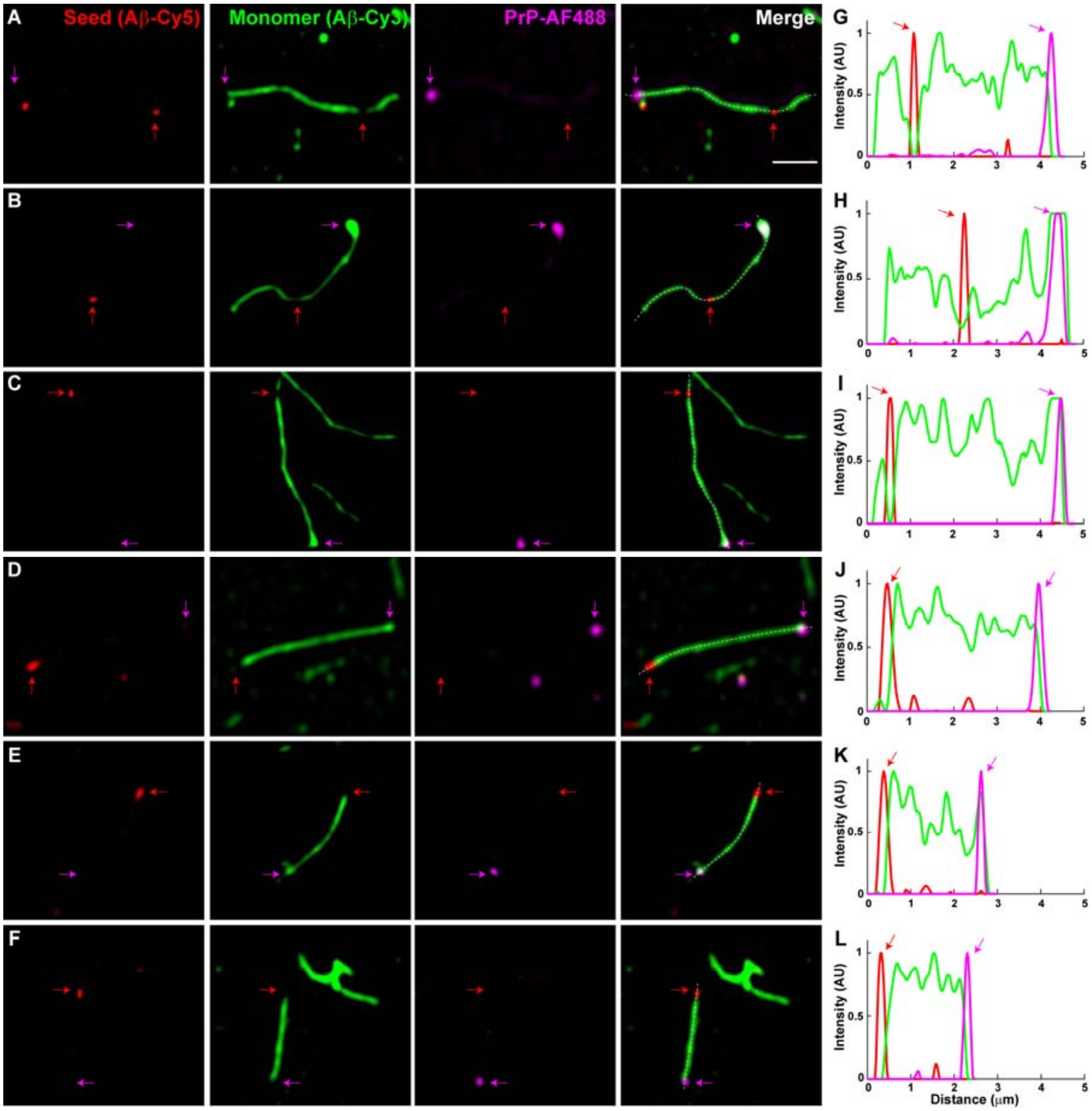


**Supplementary Fig. S3. PrP binds to fast-growing end of Aβ fibrils.** Preformed seeds composed of Aβ-Cy5 (red) were first incubated for 24 hrs with monomeric Aβ-Cy3 (green), to allow extension of the seeds at both ends. PrP-AF488 (magenta) was then added for 30 min, and the fibrils were imaged by three-color SIM. Panels **A-F** show six different microscopic fields containing individual Aβ fibrils, with each field imaged separately for Cy5 (Aβ seed), Cy3 (Aβ monomer), and AF488 (PrP), and the rightmost panel showing a merge of the three colors. The magenta arrow indicates PrP bound to the fast-growing end of a single fibril, represented by the long green extension from a red seed. The red arrow indicates the position of the seed. Scale bar in A (rightmost panel) is 1 μm. Panels **G-L** show fluorescence intensity profiles (in arbitrary units, AU) measured along the length of the fibrils, indicated by the white dotted lines in A-F (rightmost panels). Magenta and red arrows indicate peaks corresponding to the positions of the PrP and the seed, respectively.

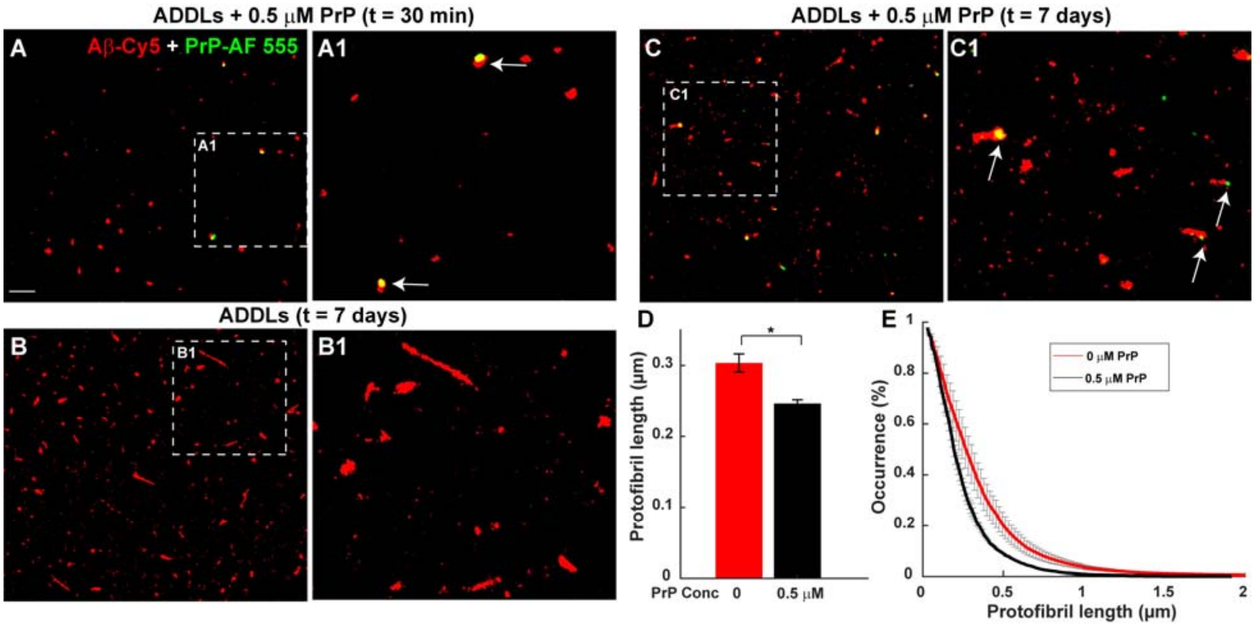


**Supplementary Fig. S4. PrP inhibits growth of protofibrils, and binds to one end of protofibrils.** **(A)** Dual color dSTORM image of ADDLs prepared from Aβ-Cy5 (20 μM, red) that were incubated for 30 min with 0.5 μM PrP-AF555 (green). Panel A1 shows the boxed area in A at higher magnification. **(B, C)** dSTORM images of protofibrils formed by incubation of ADDLs for 7 days in absence **(B)** or presence **(C)** of 0.5 μM PrP-AF555. Panels B1 and C1 show boxed areas in B and C, respectively, at higher magnification. Arrows in C1 indicate the localization of PrP at the ends of the protofibrils. **(D)** Bars show the mean lengths of protofibrils formed in the presence of 0 and 0.5 μM PrP**.** Data represent mean ± S.E. * $P<0.05$ (Student's t-test). **(E)** Cumulative distributions of protofibril lengths in the presence of 0 and 0.5 μM PrP.

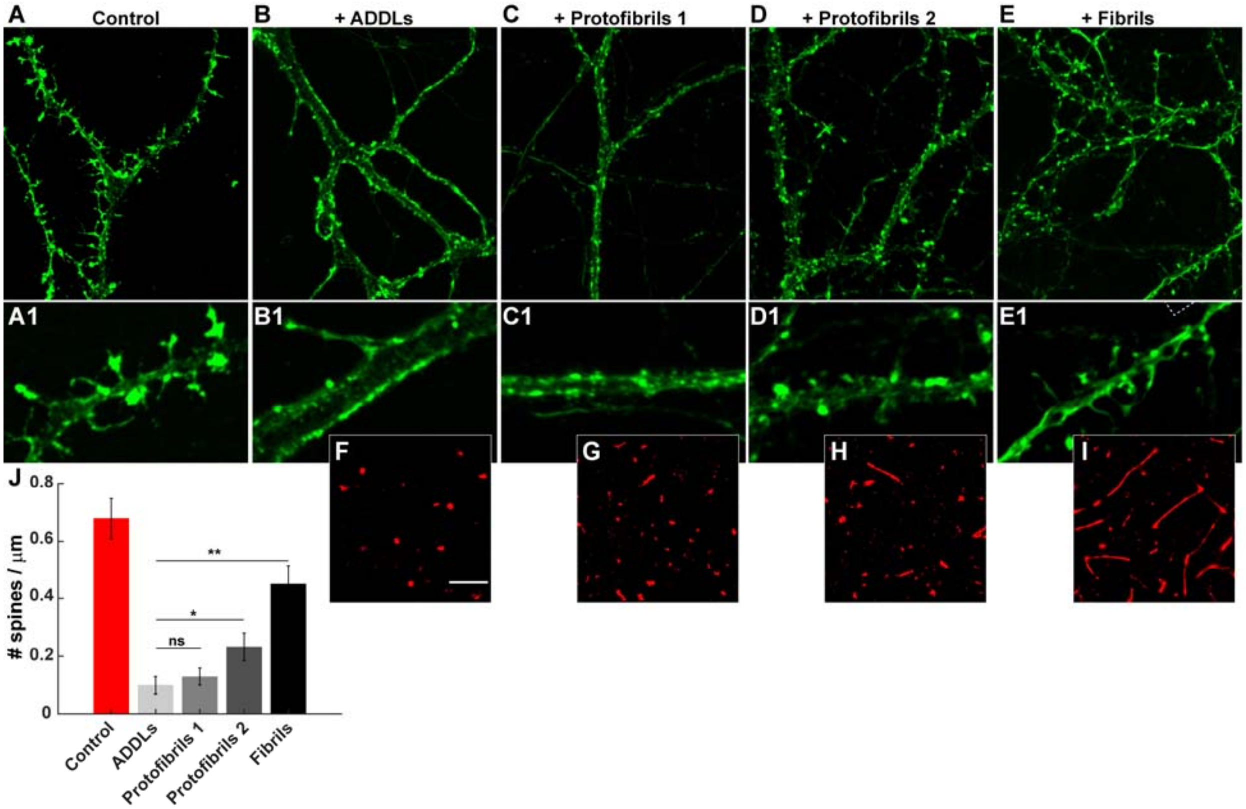


**Supplementary Fig. S5. Neurotoxicity assay of ADDLs, protofibrils, and fibrils.** Primary hippocampal neurons were untreated (Control) **(A)**, or were treated with with 500 nM (monomer-equivalent concentration) of ADDLs **(B)**, protofibril preparation 1 (incubated for 3 days) **(C)**, protofibril preparation 2 (incubated for 7 days) **(D)**, or mature fibrils **(E)**. Neurons were fixed after 24 hr of treatment and stained with Alexa 488-labeled phalloidin to visualize dendritic spines. Panels A1-E1 show boxed areas in A-E, respectively, at higher magnification. Panels **F-I** show dSTORM images of each Aβ preparation. **(J)** Quantitation of spine number per µm. Data are presented as mean± S.E. *P<0.05 and **P<0.01 (Student's t-test).

**REFERENCES**


1 Bove-Fenderson, E., Urano, R., Straub, J. E. & Harris, D. A. Cellular prion protein targets amyloid-beta fibril ends via its C-terminal domain to prevent elongation. *The Journal of biological chemistry* **292**, 16858-16871, doi:10.1074/jbc.M117.789990 (2017).

2 Klein, W. L. Abeta toxicity in Alzheimer's disease: globular oligomers (ADDLs) as new vaccine and drug targets. *Neurochemistry international* **41**, 345-352 (2002).

3 Stine, W. B., Jungbauer, L., Yu, C. & LaDu, M. J. Preparing synthetic Abeta in different aggregation states. *Methods in molecular biology (Clifton, N.J.)* **670**, 13-32, doi:10.1007/978-1-60761-744-0_2 (2011).

4 Nicoll, A. J. *et al.* Amyloid-beta nanotubes are associated with prion protein-dependent synaptotoxicity. *Nature communications* **4**, 2416, doi:10.1038/ncomms3416 (2013).

5 Cohen, S. I. *et al.* Proliferation of amyloid-beta42 aggregates occurs through a secondary nucleation mechanism. *Proceedings of the National Academy of Sciences of the United States of America* **110**, 9758-9763, doi:10.1073/pnas.1218402110 (2013).

6 Hellstrand, E., Boland, B., Walsh, D. M. & Linse, S. Amyloid beta-protein aggregation produces highly reproducible kinetic data and occurs by a two-phase process. *ACS chemical neuroscience* **1**, 13-18, doi:10.1021/cn900015v (2010).

7 Kaech, S. & Banker, G. Culturing hippocampal neurons. *Nature protocols* **1**, 2406-2415, doi:10.1038/nprot.2006.356 (2006).

8 Fang, C., Imberdis, T., Garza, M. C., Wille, H. & Harris, D. A. A Neuronal Culture System to Detect Prion Synaptotoxicity. *PLoS pathogens* **12**, e1005623, doi:10.1371/journal.ppat.1005623 (2016).
